# Are there Physical Linkages between Genes that have Synergistic Fitness Effects?

**DOI:** 10.1101/2020.03.23.004630

**Authors:** Juliet Byrnes, John Murray, Mark M. Tanaka, Ben Goldys, Antony Bellanto, Luis Cayetano, William Sherwin

## Abstract

Many of the effects on fitness in population genetics are due not to single locations in the genome, but to the interaction of genetic variants at multiple locations in the genome. Of particular interest are ‘completely epistatic’ interactions, where a combination of genetic variants is required to produce an effect, and the effect cannot occur with any other combination. In diploids, epistasis is strongly connected to meiotic recombination, a process which can both assemble and destroy beneficial combinations of genetic variants. Additionally, epistatic interactions can be hard to detect in empirical studies, and mathematical models of epistasis and recombination are challenging to analyse, so despite their ubiquity epistatic interactions are regularly not considered. As a result, there is little consensus on when high levels of recombination might be expected, or how strongly recombination affects beneficial or deleterious fitness effects controlled by epistatic interactions. We address this question by conducting a meta-analysis and simulations. The meta-analysis used data drawn and curated from *Drosophila melanogaster* studies in Flybase. We extracted studies relating genetic combinations and phenotypically detectable effects on fitness, then analysed the relationship between the rate of recombination and effect on fitness with a statistical model. We also ran simulations under a two-locus Wright-Fisher model with recombination and epistatic selection. The results of both approaches indicated a tendency for genetic combinations with an epistatic effect on fitness to occur in an environment of reduced meiotic recombination. Two possible explanations for this are that the variants controlling such interactions are selected for in regions where there is little recombination, or that such interactions lead to selection for lower rates of recombination in the regions where those variants appear.

## Introduction

Theories of the interaction of meiotic recombination and selection have had a long and tumultuous history, with almost a century of research dedicated to understanding how a system as reproductively costly as recombination could have evolved and why it is pervasive in so many species (Otto and Lenormand 2002). Recombination creates but also destroys ‘haplotypes’, DNA molecules carrying combinations of alleles at different loci. Selection is also complex, especially when based on allelic variation at two or more loci in haplotypes. Of particular importance is epistasis, a multilocus selection effect in which alleles at different loci interact to produce an effect on fitness, either synergistically or antagonistically, that neither could produce in the absence of the other. Synergistic epistasis means that if each locus has one allele with the same type of fitness effect (i.e. both are either beneficial or deleterious), the combination of those alleles will have a stronger effect on fitness than effects of both the alleles if they are assumed to act independently. In contrast, under antagonistic epistasis, that combination of alleles will have a weaker effect. When neither allele has a fitness effect by itself, with the effect only appearing when both alleles are present, this is known as complete epistasis. If we arbitrarily label the alleles at each locus ‘0’ and ‘1’, then we call the genetic types (or ‘haplotypes’) with a ‘0’ at both loci or a ‘1’ at both loci the ‘coupling haplotypes’, and the remaining two haplotypes the ‘repulsion haplotypes’ [TABLE 1]. Henceforth we consider the coupling haplotype with ‘0’s at both loci to be under selection that is completely epistatic (no single locus selection effects occur) and label it as the ‘beneficial haplotype’ or the ‘deleterious haplotype’ depending on the direction of the selection effect. We refer to the haplotype with ‘1’s at both loci as the ‘non-beneficial coupling haplotype’ or the ‘non-deleterious coupling haplotype’ also depending on the direction of the same selection effect, meaning that it has the same fitness value as the repulsion haplotypes. Epistasis in its various forms can speed up or slow down selection, can induce selection plateaus (Moore and Williams 2015), and is regularly considered as a possible driver of the evolution of recombination rates.

**TABLE 1:** Table describing the names of the multilocus haplotypes we use in this paper. These are consistent with their usual names in the population genetics literature.

|  |  | Locus 1 |  |
| --- | --- | --- | --- |
|  |  | Allele 0 | Allele 1 |
| Locus 2 | Allele 0 | (0,0) Coupling, beneficial or deleterious | (0,1) Repulsion |
|  | Allele 1 | (1,0) Repulsion | (1,1) Coupling, non-beneficial or non-deleterious |

Possible explanations for the prevalence of recombination are broadly divided into three categories, with epistasis often playing a prominent role (Otto 2009, Hartfield and Keightley 2012).

1. It has been proposed that recombination 1) prevents beneficial mutations at separate loci from competing with one another and 2) allows the creation of chromosomes with no deleterious mutations from two chromosomes that each have a deleterious mutation at different loci. These phenomena are termed ‘clonal interference’ (Hill and Robertson 1966) and ‘Muller’s Ratchet’ respectively (Muller 1932, Felsenstein 1974). A recurring theme in the recombination literature deals with the speed of production of double mutants - i.e. a haplotype containing novel mutations at two different loci - particularly if those mutations are beneficial. An extensive body of theoretical literature has shown that higher rates of recombination cause a double mutant to occur much earlier in an evolutionary history than otherwise (Barton 1995, Suzuki 1997, Christiansen *et al*. 1998, Weissman *et al*. 2010).
2. Another argument proposes that recombination makes a species more successful under fluctuating selection, such as by allowing rapid production of novel, possibly adaptive haplotypes when a species is co-evolving with its parasite (Kondrashov & Yampolsky 1996, Peters and Lively 1999). Therefore some authors have considered models with fluctuating levels of selection (Sasaki and Iwasa 1986), epistasis (Suzuki 1997, Weissmann *et al*. 2010) or fluctuating selection *and* epistasis (Peters and Lively 1997, Otto and Michalakis 1998, Gandon and Otto 2007).
3. A third proposal is that recombination increases the effectiveness of selection in purging deleterious mutations when they are common and act synergistically (Kondrashov 1988, Gandon and Otto 2007). Empirical explorations of this by Fry *et al*. (2004) and Rosa *et al*. (2005) found evidence of synergistic epistasis in *Drosophila melanogaster*, though Rosa *et al*. contend that this is too occasional to explain the origin or maintenance of meiotic recombination.

The situation is complicated by the fact that early models of recombination, which did not include genetic drift, showed no benefit to fitness of recombination (Felsenstein 1974; Otto and Barton 2001), despite the fact that recombination is ubiquitous and highly variable at every scale (Stapley *et al*. 2017). Later stochastic models of recombination, which do include genetic drift, pose strong mathematical difficulties, though there has been some progress in the mathematical analysis of these models (Mano 2013, Jenkins and Song 2015, Esser *et al*. 2016). An important simplification is to restrict consideration to populations that are at ‘linkage equilibrium’, i.e. the proportion of a haplotype(*i, j*) in the population is equal to the proportion of haplotypes possessing the *i* allele at the first locus times the proportion of haplotypes possessing the allele at the second locus (*p_ij_* = *p_i_p_j_*). When this occurs and there is no selection, evolution at each locus can be modelled independently (Durrett 2008, Mano 2013). As a consequence, the quantity *P_ij_* − *P_i_P_j_* is useful in modelling recombination, and is known as a measure of ‘linkage disequilibrium’.

However, recombination can have positive, negative or equivocal effects. For example, recombination can clearly be a disadvantage due to breaking up selectively advantageous haplotypes. Therefore, the faster production of double mutants in the presence of recombination would be offset by a relative increase in the time taken for the double mutant to reach fixation in the population when compared to the non-recombining case (Barton 1995, Suzuki 1997, Christiansen *et al*. 1998). Additionally, Weissman *et al*. (2010) contend that there is an optimal rate of recombination that causes the appearance of the first double mutant to occur much earlier if the alleles experience epistatic selection – but when some ‘critical’ recombination rate is exceeded, the first double mutant will take an exponentially longer time to first occur. However, these models have also been criticised for being too restrictive for an underlying biological mechanism to have evolved in the first place (Baumgardner *et al*. 2013) or for requiring a parameter space that is too restrictive to allow persistence (Otto 2009).

It can be seen from the above that plausibly there are both advantages and disadvantages for recombination between epistatically interacting loci. The fitness effects epistasis could have on ecological and genetic processes mean that there are important theoretical and empirical implications from understanding the relationship between epistasis and recombination. We therefore investigated the relationship between recombination, epistasis, selection and drift in two ways: 1) via a meta-analysis using Flybase and 2) via simulations of small randomly mating populations under epistatic selection at two recombining loci. Specifically, we address the following three questions about pairs of loci that interact synergistically to have an effect on fitness:

1. If the interaction is deleterious, do the loci recombine more often than the average pair of loci, as proposed by Kondrashov? Or will they have a lower rate of recombination, as suggested by the body of evidence in Otto and Lenormand’s review (2002)?
2. If the interaction is beneficial, do the loci recombine more often than the average pair of loci, as suggested by clonal interference and Muller’s Ratchet? Or will they have a lower rate of recombination?
3. the interaction is beneficial, do the loci have an optimal rate of recombination, as suggested by Weissman *et al*. (2010)?

## Methods

### Flybase Meta-analysis: Data on Interactions and Recombination

We were interested in whether the fitness of a phenotype determined by pairs of genetic loci had an effect on the amount of recombination between those loci. We reviewed a number of different databases with genetic data on various species (The Arabidopsis Information Resource (Berardini *et al*. 2004), MaizeGDB (Andorf *et al*. 2015), ZFIN (Howe *et al*. 2013), Flybase (Gramates *et al*. 2016)) to determine suitability of use in conducting this investigation. Our criteria for selection were: 1) the database links alleles to observed phenotypes in the study species; 2) the expression of several observed phenotypes depends on the presence of particular alleles at 2 or more loci; and 3) the alleles at these loci were naturally occurring and the genetic loci involved had been mapped genetically, so that we could determine recombination rates. These criteria were satisfied only by Flybase, a database containing an extensive set of genetic and phenotype data on *D. melanogaster*.

First, we identified the pairs of genetic loci that determined a phenotype and whether there was epistasis between them; second, we estimated the recombination rates between these genetic loci based on genetic map data; thirdly and finally, we fitted a regression model to analyse the data with recombination rates as the response and ‘epistasis class’ as the predictor variables, where the ‘epistasis class’ of a locus pair is defined as the effect of that locus pair on the genotype of a fly. Epistasis class was determined by extracting data from Flybase on phenotypes, the genetic loci that determine that phenotype and the particular phenotype information associated with certain combinations of alleles at those loci. We considered phenotypes determined by pairs of loci only, and sorted them into three epistasis classes; those where the genetic loci had 1) a phenotypically discernible interaction, with an impact on fitness (abbreviation PDFI); 2) a phenotypically discernible interaction, which may or may not have an impact on fitness (abbreviation PDNF); or 3) with no phenotypically discernible interaction (abbreviation NDI), i.e. the genetic loci acted on the phenotype independently but not epistatically. We processed the data by extracting the results of studies on the effect of various genetic loci on phenotypes, excluding pairs involving alleles which did not occur naturally, and stratified them into types of interaction.

We searched for haplotypes consisting of alleles at two or more loci, and whether these haplotypes had any epistatic effects on fitness. We used a genetics interaction file previously computed by Flybase on allele interactions and the phenotypes associated with them. We refer to this as [GENE_INT]. We also used the Flybase SQL database for the Flybase meta-analysis. The files used are the February 2017 release. The table in [GENE_INT] contains 4 columns: 1) the symbol (a human readable name) of an allele in the interaction; 2) the unique Flybase reference of the allele; 3) the description of the type of interaction and all genetic loci and alleles involved; and 4) the unique Flybase reference of the study which determined the genetic loci involved in the phenotype. An example of the tabulation in Flybase is given in [TABLE 2].

**TABLE 2:**
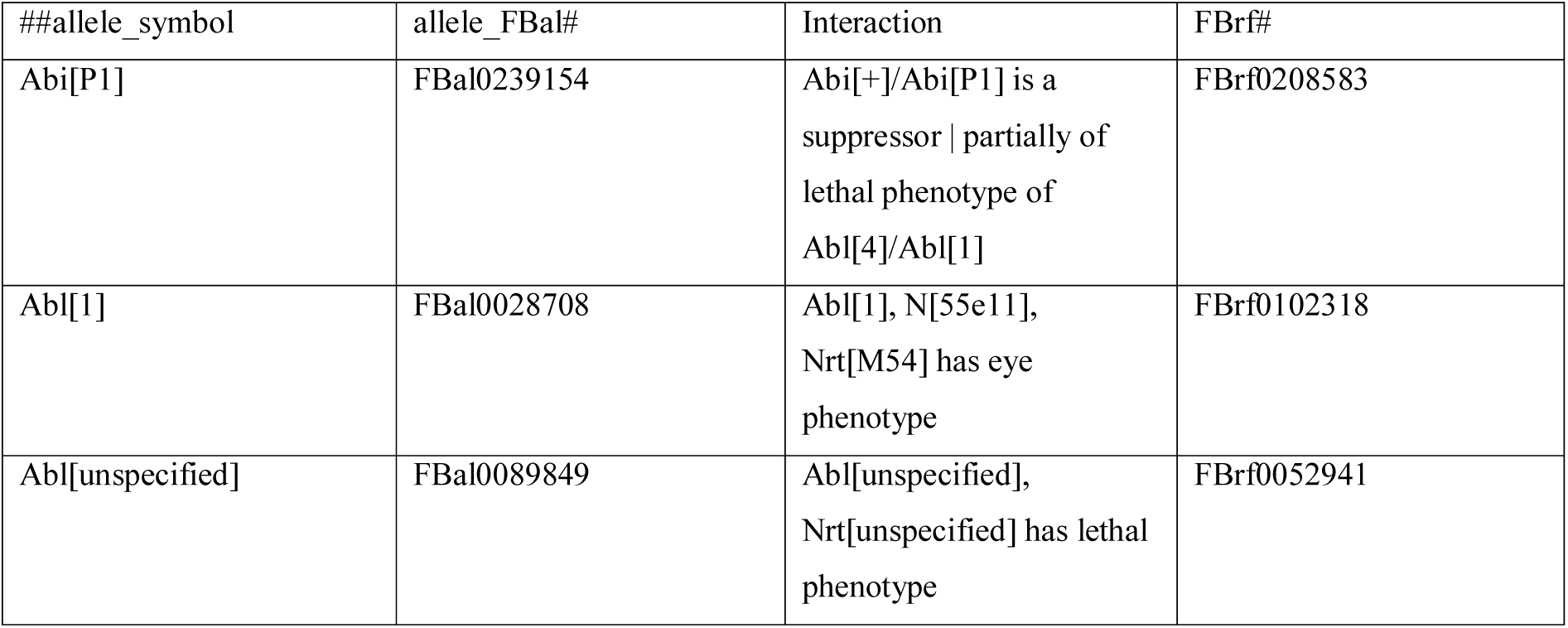
An example of what a phenotype-gene association looks like when obtained from Flybase. The first column gives a locus name followed by the allele name in square brackets. The second column gives the unique Flybase reference for this allele. The third column gives the names of all alleles and the loci they exist on involved in the interaction as well as the type of interaction. In the first row, third column, there is a lethal interaction between the loci ‘Abi’ and ‘Abl’. In the second row, third column, there is a non-fitness interaction between three loci; ‘Abl’, ‘N’ and ‘Nrt’. In the third row, third column, there is a lethal interaction between the loci ‘Abl’ and ‘Nrt’.

Each data entry contains a phenotype and the alleles associated with it, so there may be multiple data entries which involve the same sets of loci. Our search method identified the names of the genetic loci involved in each data entry. Details of this search method can be found in the supplement [Code Supplement, Part I: Input and Search Patterns]. There were 98087 entries for *D. melanogaster* locus pairs. We identified 81265 phenotypes determined by sets of loci containing two loci, and 9585 containing three or more. We removed phenotypes involving more than three loci for simplicity. In practice, 6754 out of the 9585 (70.5%) cases where there was a three-way interaction also contained a two-way interaction. We then standardised the names of the genetic loci by using a SQL query to locate their unique Flybase references and name by symbol. This was necessary because a single locus can have multiple different symbolic names in the Flybase database, but will always have a unique Flybase reference ID. Because we were only interested in alleles which occur in natural populations or spontaneously in cultivated populations, we used the SQL database to extract the list of mutagens associated with each allele. We then removed any entries where a genotype had an allele with a mutagen listed other than ‘natural population’. Of the original 98087 locus pairs, only 5346 locus pairs were associated with natural mutations.

We then classified these 5346 locus pair entries into three ‘epistasis classes’ based on their effects on the phenotype. If an entry contained the terms ‘non-suppressor’, ‘non-enhancer’, ‘non-enhanceable’ or ‘non-suppressible’, this meant that the allele at one of the loci in the data entry had no discernible effect on the expression of a phenotype associated with the allele at the other locus, i.e.: they acted on the phenotype independently but not epistatically. We were able to detect these in entries by reducing them to the stems ‘non-enh’ or ‘non-suppr’. Furthermore, the fitness effects recorded for the data entries consisted of four types. They were: 1) lethality: where flies possessing the genotype had a higher death rate than those without the genotype; 2) sterility: where flies possessing the genotype had fewer offspring than those without the genotype; 3) fertility: where flies possessing the genotype had *more* offspring than those without the genotype; and 4) viability: where flies possessing the genotype were more likely to survive than those without the genotype.

We classified these effects into epistasis classes according to the following protocol:

1. PDFI: phenotypically detectable fitness interactions: the alleles at the genetic loci interacted and this had a discernible impact on fitness in the form of survival or fertility. These did not contain the stems ‘non-enh’, ‘non-suppr’, but did contain the terms ‘lethal’, ‘fertile’, ‘sterile’, or ‘viable’.
2. PDNF: phenotypically detectable interactions with no known impact on fitness: the alleles interacted, but their interactions had no visible impact on fitness. These did not contain the word stems ‘non-enh’, ‘non-suppr’, and did not contain the terms ‘lethal’, ‘fertile’, ‘sterile’ or ‘viable’.
3. NDIs: No phenotypically discernible interaction: the alleles at the genetic loci did not interact. In the Flybase data, these all contained the abbreviated stems ‘non-enh’ or ‘non-suppr’, indicating that the alleles at each loci acted only independently on the phenotype and not epistatically (the allele at one locus neither enhanced nor suppressed the phenotypic effect of the allele at the other locus).

Where a single pair of loci had different allele combinations appearing in the data that would allow entry to more than one of the three categories above, we used the following decision method [FIGURE 3]:

1. If any observed combinations of alleles at the locus pair were classified as PDFI, the locus pair was classified as PDFI.
2. If any observed combinations of alleles at the locus pair were classified as PDNF, and none were classified as PDFI, the locus pair was classified as PDNF.
3. If no observed combinations of alleles at the locus pair were classified as PDFI or PDNF, the locus pair was classified as NDI.

As a result of this process, many (5076) duplicate locus pair entries were removed, leaving a total of 270 unique locus pairs. We classified 39 locus pairs as PDFI, 184 locus pairs as PDNF, and 47 locus pairs as NDI.

**FIGURE 3:**
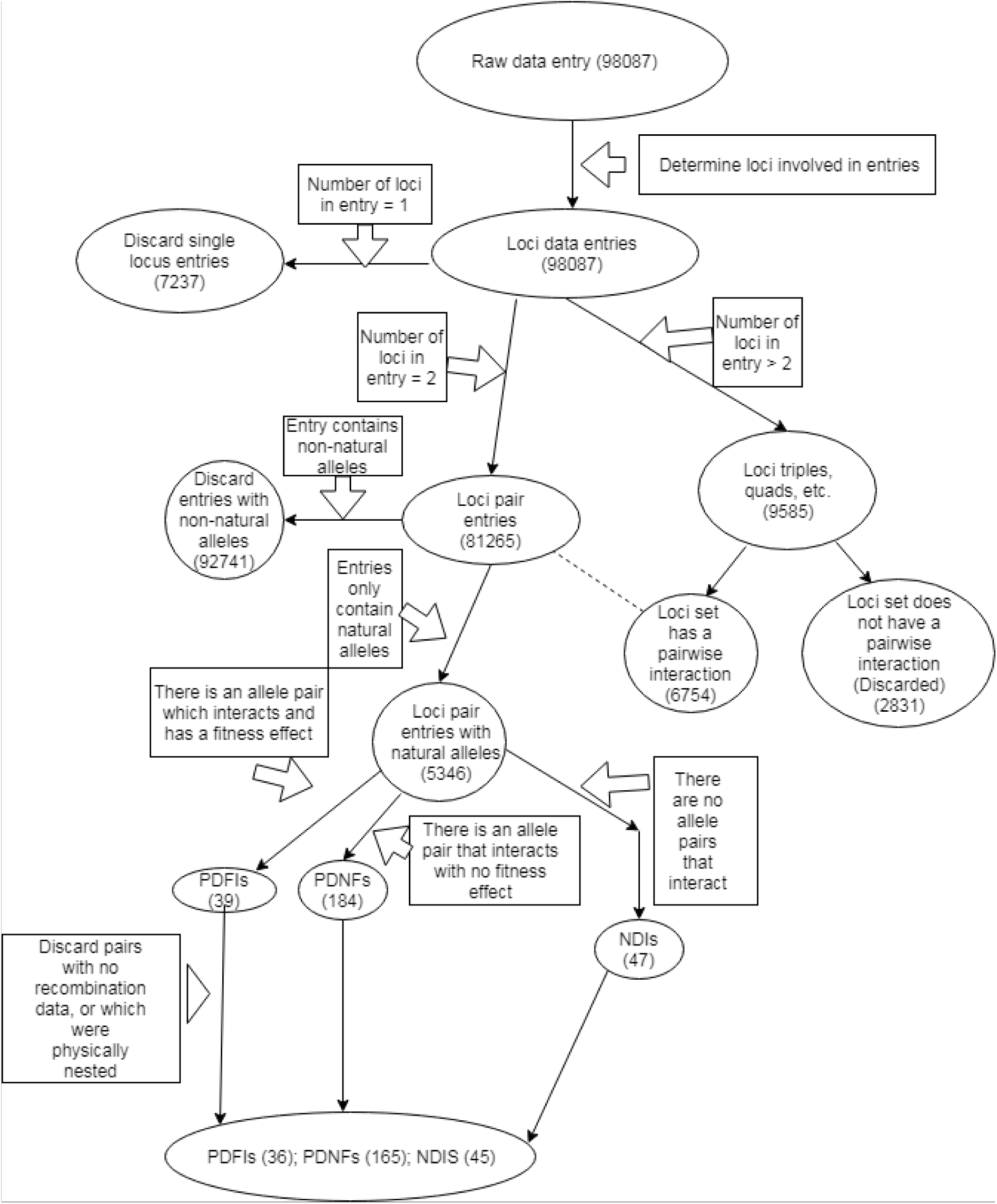
A flowchart describing the data extraction process and the number of data points at each step. NDIs are cases where both loci affect the phenotype, but there is no epistatic interaction. The bottom discard step applies to all classes.

We used the genetic map data from the precomputed Flybase file to filter for locus pairs that were on the same chromosome, because meiotic homologous recombination only occurs between loci lying on the same chromosome. The genetic map data also gave the average rate of recombination between each locus and a designated end of the chromosome (telomere). These data may have been determined by experiment or computed using information from nearby loci, depending on the locus of interest. The unit for recombination is centimorgans (cM), where 1 cM indicates that the average number of recombinations expected in a generation is 0.01. We then use this to approximate the probability of recombination between two loci – if two loci are 1 cM apart, the probability of a recombination occurring in a single generation between the two loci is 0.01. To reflect the biological reality of the maximum possible observed recombination probability in diploid organisms, we truncated the probabilities of recombination at 0.5. We discarded locus pairs where recombination data were not available for one or more of the loci involved. This resulted in the removal of 21 data points – 2 (out of 47) NDIs and 19 (out of 184) PDNFs. Finally, we removed interactions where one locus was nested inside another (as loci can span several megabases). This removed 3 (out of 39) PDFIs. In the final extracted dataset there were 36 PDFIs, 165 PDNFs and 45 NDIs [TABLE 7]. 53 locus pairs were on the X chromosome, 100 pairs were on chromosome 2, and 93 pairs were on chromosome 3. There were no locus pairs located on chromosome 4. The final dataset contained the recombination rate between each pair of loci (*c*), the epistasis class of the two loci, and the chromosome on which the pair of loci were located (which we will call ‘chromosomal identity’). Out of a total of 246 locus pairs, there were 19 locus pairs (7.7%) with the minimum recombination rate of *c =* 0 and 51 locus pairs (20.7%) with the maximum recombination rate of *c =* 0.5.

### Flybase Meta-analysis: Regression Model to Test for Effects of Epistasis Class and Chromosome Number on Recombination

For our study, we wished to know whether the epistasis class of a locus pair drove the probability of recombination between the two loci. We also needed to control for chromosomal identity due to the large variance in recombination rates between chromosomes. We needed a model that could deal with data having a large number of points on 0 and 0.5 plus a spread of values in between. Therefore we analysed the extracted dataset using a partially stratified zero-and-one-inflated beta regression model (Ospina and Ferrari 2012). The zero-and-one-inflated beta regression model contains sub-units which we will call ‘components’, and each component contains one response variable, a set of explanatory variables, a link function, and parameters associated with each explanatory variable in the generalised linear model framework. Therefore when modelling the recombination rate as a response of the zero-and-one-inflated beta regression model the response variables of the four sub-model components are:

1. the mean of the intermediate recombination rates (i.e. recombination rates that were not 0 or 0.5);
2. the precision of the intermediate recombination rates, defined by precision = [mean(1-mean) / variance] - 1;
3. relative probability that *c =* 0 rather than 0 < *c <* 0.5; and
4. the relative probability that *c =* 0.5 rather than 0 < *c* < 0.5.

The first and second components model the recombination rates that are not at the extremes of 0 and 0.5 (the Intermediate Recombination Rates) according to a beta distribution. The precision variable describes the relationship between the variance and the mean and is allowed to vary with different explanatory variables. The third component models the probability of the recombination being 0 divided by the probability of the recombination rate lying in the intermediate range of 0 < c < 0.5 (the Odds of Minimum Recombination). The fourth component models the probability of the recombination rate being 1 divided by the probability of the recombination rate lying in the intermediate range of 0 < c < 0.5 (the Odds of Maximum Recombination).

The link functions for each component response variable are given in [TABLE 4] Each component sub-model shares the same set of explanatory variables. These were epistasis class, chromosomal identity and the interaction of epistasis class and chromosomal identity. However, each sub-model can - and in practice, will - have different parameter fits for each explanatory variable.

**TABLE 4:**
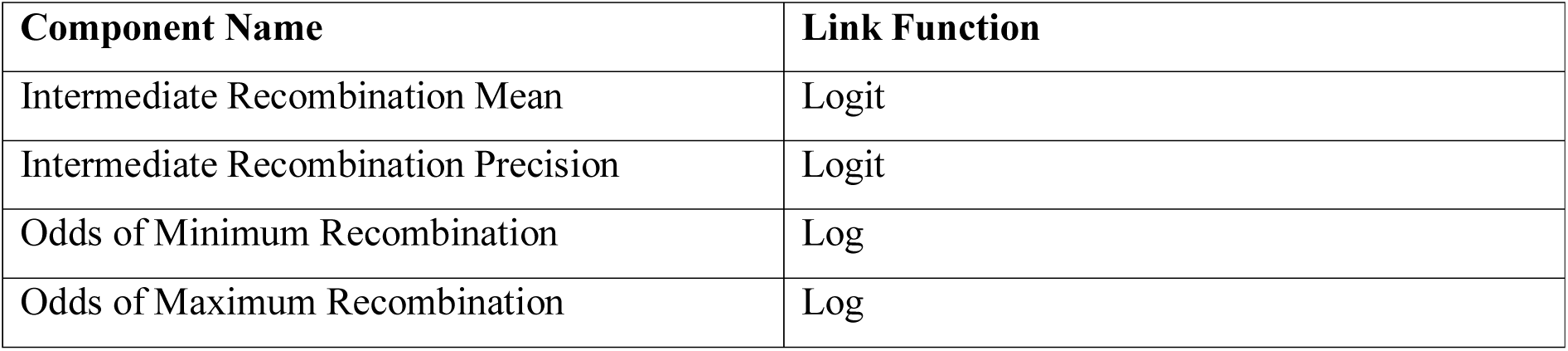
The link functions for each component of the zero-and-one-inflated beta regression model. Each component submodel is of the form: *link(responds)* = ∑*_i_ a_i_x_i_* where the *x_i_*’s are the explanatory variables, the *a_i_*’s are the parameters of the sub-model component, and is the index of the covariates included in the component.

Epistasis class was coded as two binary variables. The first epistasis variable was 1 if the epistasis class was PDFI, and 0 otherwise; we labelled this variable as ‘Fitness’. The second variable took the value 1 if the epistasis class was PDNF *or* PDFI, and 0 otherwise; we labelled this variable ‘Interaction’. If both variables were zero, the epistasis class was NDI. There were also two binary variables coding chromosomal identity. The first variable took the value 1 if the locus pair was on chromosome 2 and 0 otherwise; we labelled this variable ‘Chrom2’. The second variable took the value 1 if the locus pair was on chromosome 3 and 0 otherwise; we labelled this variable ‘Chrom3’. If both of these variables were 0, the locus pair was on chromosome 1 (the X chromosome). The parameter coding is shown in [TABLE 5].

**TABLE 5:**
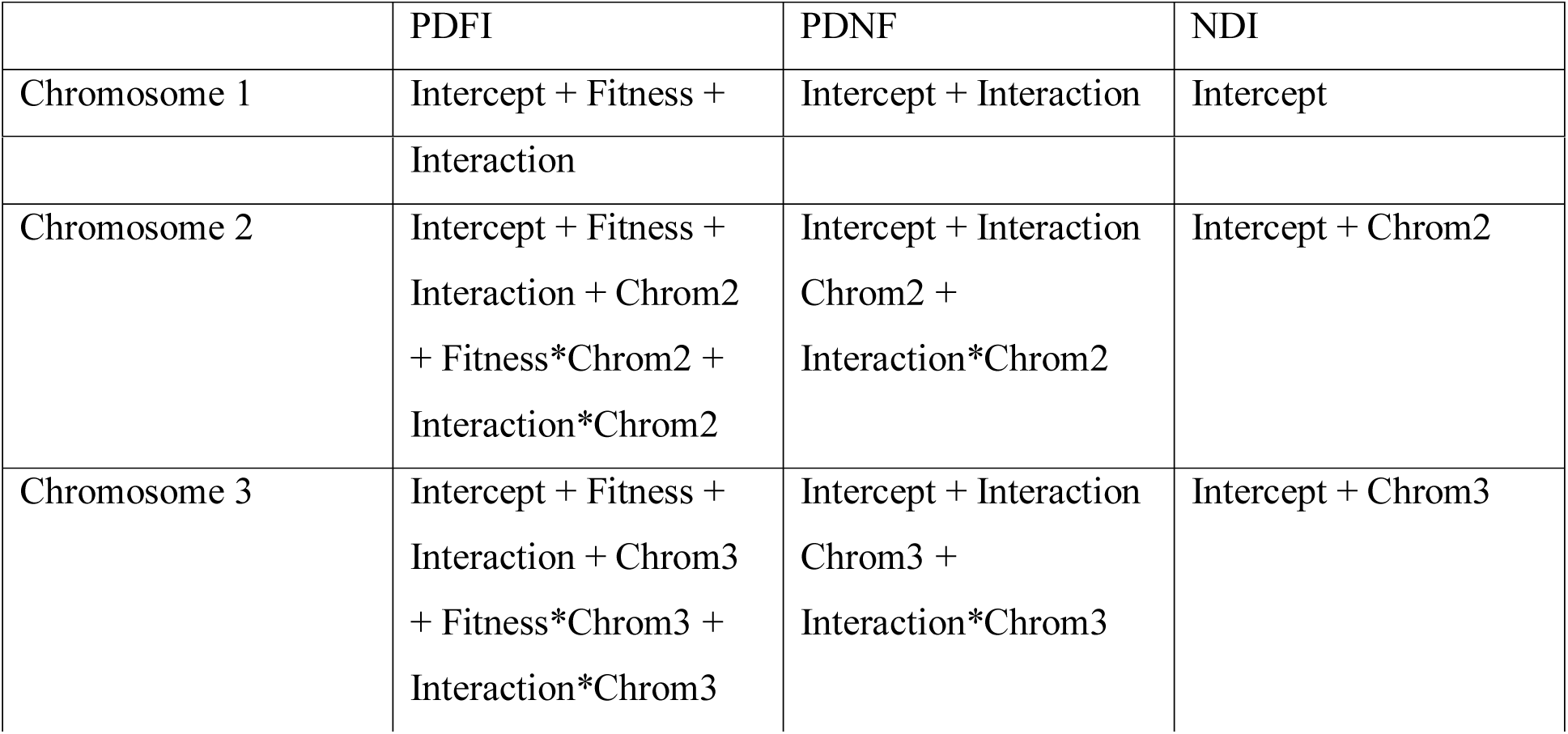
Coding of the variable parameters for the nine different categories for a component sub-model containing all the explanatory variables stated above (epistasis class, chromosomal identity and their interactions). The ‘Intercept’ parameter is *not* a true baseline, because the variables are categorical. Four components result in a total of 32 parameters to fit if all explanatory variables are included in all the component sub-models. The parameter for a locus pair on chromosome 1 in the NDI class is coded as the intercept. Variables with a ‘*’ between them indicate interactions of those two variables.

We could not include all explanatory variables in every sub-model because our sample size was very small (246) compared to the number of parameters described by having two categorical variables with three levels (epistasis class and chromosomal identity) and the interaction of those variables (8 parameters), and having four model components (32 parameters in total). We therefore required a model selection procedure. We used a generalized Akaike information criterion (GAIC) provided by the R package GAMLSS (Rigby and Stasinopoulos 2005) to simultaneously perform model selection and fit the parameters of the model. The GAIC model selection procedure excludes explanatory variables from each sub-model if they do not contribute substantially to the model fit.

We provide a brief explanation of the interpretation of the parameters that describe the relative probabilities of the minimum and maximum recombination rates. We assign each locus pair to a category, which is the particular combination of chromosome and epistasis class to which it belongs [TABLE 7]. For instance, a locus pair on Chromosome 3 and in epistasis class NDI would be in category (NDI, 3). We consider the component (4) above, which describes the relative probability that *c =* 0.5 rather than 0 < *c* < 0.5. The response variable of this component sub-model is the log of the probability that pairs of loci in a particular category freely recombine (at the maximum recombination value, *c* = 0.5) divided by the probability that pairs of loci in the same category recombine at an intermediate rate (0 < *c <* 0.5), i.e.:

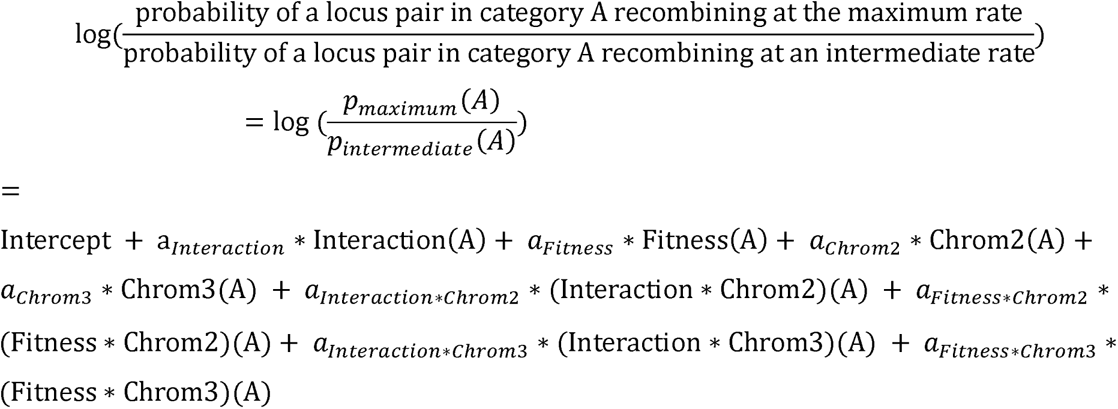

Note that if we consider the category to be the (NDI, 1) category, we get:

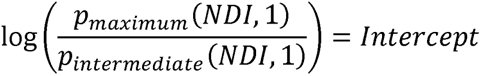

Similarly, if we consider category to be the (PDNF, 3) category, we get:

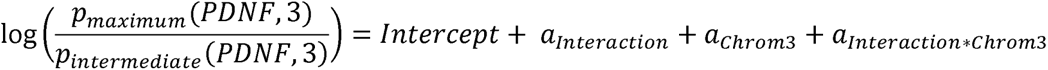

Each parameter or some sum of parameters (excluding the intercept) therefore has an interpretation as the log of an odds ratio. For instance:

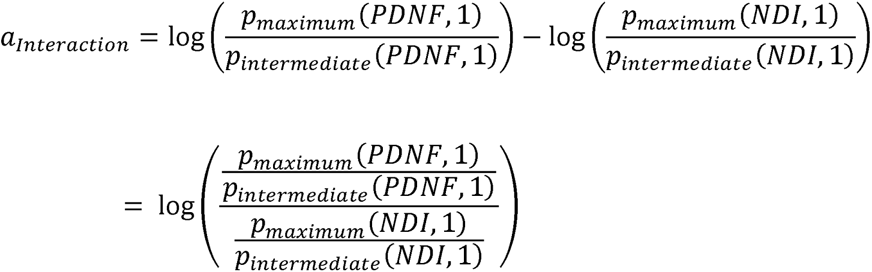

So 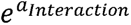 is a ratio of odds. The numerator of this ratio is the odds of a locus pair in the (PDNF,1) category recombining at the maximum rate over recombining at an intermediate rate. The denominator of this ratio is the odds of a locus pair in the (NDI,1) category recombining at the maximum rate over recombining at an intermediate rate. If the ratio is greater than 1 (>1), this means that the odds of a locus pair in the (PDNF,1) category recombining at the maximum rate over recombining at an intermediate rate is greater than the corresponding odds for a locus pair in the (NDI,1) category.

The conclusion is the opposite if the ratio is less than 1 (<1) and the odds are equal if the ratio is equal to 1. This interpretation is why the component is referred to as the ‘Odds of Maximum Recombination Component’. To illustrate this, suppose that the probability of a locus pair in the (PDNF,1) recombining at a maximum rate is 0.2 and the probability of an intermediate rate is 0.1.

Then the odds for this is 2. Further suppose that the probability of a locus pair in the (NDI,1) class recombining at a maximum rate is 0.3 and the probability of an intermediate rate is 0.3. Then the odds is 1. Now the odds ratio is 2, and we can follow the interpretation above, but the *probability* of a maximum rate for a locus pair in the (PDNF,1) class is in fact *smaller* than the probability of a maximum rate for a locus pair in the (NDI,1) class. Some care must therefore be taken with the interpretation of results. All the arguments above are the same with 0s instead of 0.5s for the Odds of Minimum Recombination Component.

### Simulations of Epistatic Selection and Recombination

We simulated the evolution of a small population at two loci according to a single-sex Wright-Fisher haploid model with recombination and synergistic epistatic selection. The Wright-Fisher haploid model assumes a fixed population size and random mating. The next generation is determined by randomly drawing individuals from the previous generation under a multinomial distribution with probabilities determined by the proportion of the haplotype in that generation and the selection strength. With recombination, the model is modified to include a binomial element – for each new haploid individual, there is a probability ‘*c*’ of recombination while reproducing. When a recombination event occurs, another individual is drawn and the haplotype is modified by recombination between the two individuals drawn. We further modified this model by using a beneficial complete epistasis fitness scheme. This scheme did not include sex and is technically a haploid model with recombination. In general, however, haploid models provide very good approximations to diploid models of half the size in most cases and it is a convention to use haploid models where possible (Durrett 2008).

Population size was varied in multiples of 50, with the smallest containing 50 individuals and the largest containing 500 individuals. We modelled haplotypes of two biallelic loci, with haplotypes labelled as in [TABLE 1]. We set the relative fitness of the beneficial haplotype to be 1 + *s*, *where s* was allowed to vary from 0 to 0.05, and the other three haplotypes had fitness 1. We varied the recombination probability *c* from 0 to 0.025. We then investigated higher recombination probabilities with simulations in which recombination probabilities ranged from 0.01 to 0.5, but only with a single population size (200). We also considered twelve different cases for the initial genotype frequencies with different levels of linkage disequilibrium and relative proportions of the beneficial genotype in the population. The details can be found in [TABLE 6].

**TABLE 6:**
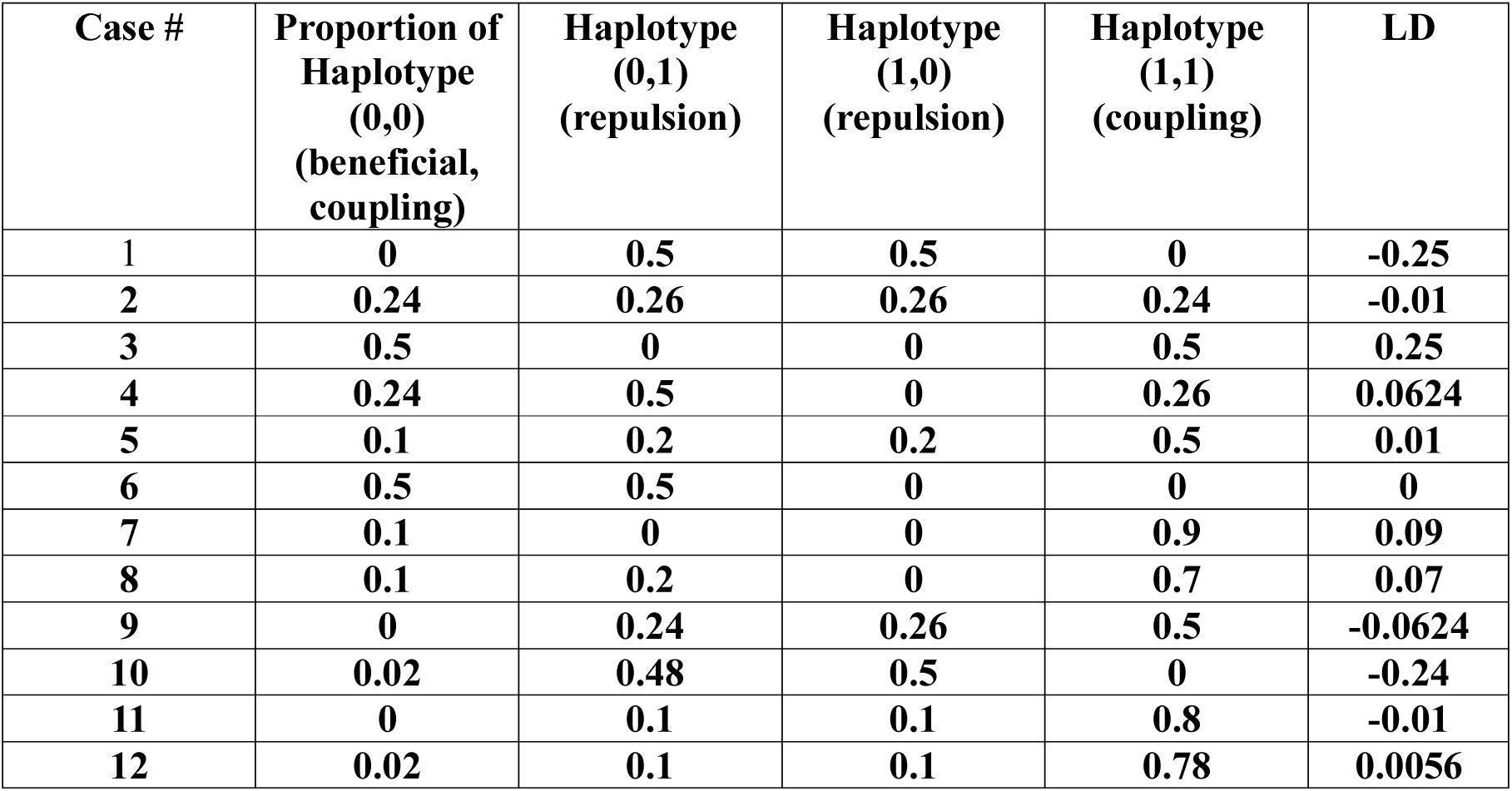
The initial haplotype proportions for each of the twelve cases, and their linkage disequilibrium, calculated by *p*_00_ − (*p*_01_ + *p*_00_)(*p*_10_ + *p*_00_)

Each simulation ran for 50 generations, with 10,000 replicates per simulation. This period of time was sufficient for several simulations to reach approximate linkage equilibrium; these patterns are reflected in our graphs [FIGURE 10, FIGURE S3 Cases] We assessed the behaviour of the model by examining the probability that beneficial haplotype becomes fixed in the population (100%) and the mean time for this fixation, if it occurred. The code for the simulations is written entirely in R and can be found in the appendix.

## Results

### Flybase Meta-analysis

The zero-and-one-inflated beta regression model appeared to be an appropriate model for the recombination data. A model where either the distribution of link function was incorrectly specified would likely display a pattern in the residuals. The residual plots for the fitted model showed no detectable trend, supporting the choice of the zero-and-one-inflated beta regression model [FIGURE S5].

There were 36 locus pairs with phenotypically detectable fitness interactions [PDFIs], 165 locus pairs with phenotypically detectable interactions without a fitness effect [PDNFs] and 45 locus pairs with no phenotypically detectable interaction [NDIs] in the final data set. (These abbreviations are defined in Methods). Out of a total of 246 locus pairs, there were 19 locus pairs (7.7%) with the minimum recombination rate of *c =* 0 and 51 locus pairs (20.7%) with the maximum recombination rate of *c =* 0.5.

All observed fitness interactions in the final dataset were lethal, i.e. the epistasis was deleterious. There was substantial variability in the locations of the locus pairs across the genome. Most pairs of loci in the extracted dataset were on chromosome 2 (see [TABLE 7]). On each chromosome the sampled loci could be quite close to a telomere or the centromere. Distance in base pairs between the paired loci ranged from zero to almost three-quarters of the chromosome [FIGURE S1]. Due to the large parameter space relative to sample size, we used a generalized Akaike information criterion (GAIC) to select the best model (see in Methods: Flybase Meta-analysis: Regression Model to Test for Effects of Epistasis Class and Chromosome Number on Recombination).

**TABLE 7:** Numbers of locus pairs in each of the nine possible categories after filtering.

|  | **PDFI** | **PDNF** | **NDI** |
| --- | --- | --- | --- |
| **Chromosome 1** | 6 | 40 | 7 |
| **Chromosome 2** | 12 | 73 | 15 |
| **Chromosome 3** | 18 | 52 | 23 |

According to the GAIC the best model was the intercept-only model for the intermediate recombination mean, intermediate recombination precisions and odds of minimum recombination components, and for the odds of maximum recombination component included the epistasis class and chromosomal identity variables without interactions ([TABLE 8], [FIGURE S2] & [TABLE 9]).

**TABLE 8:**
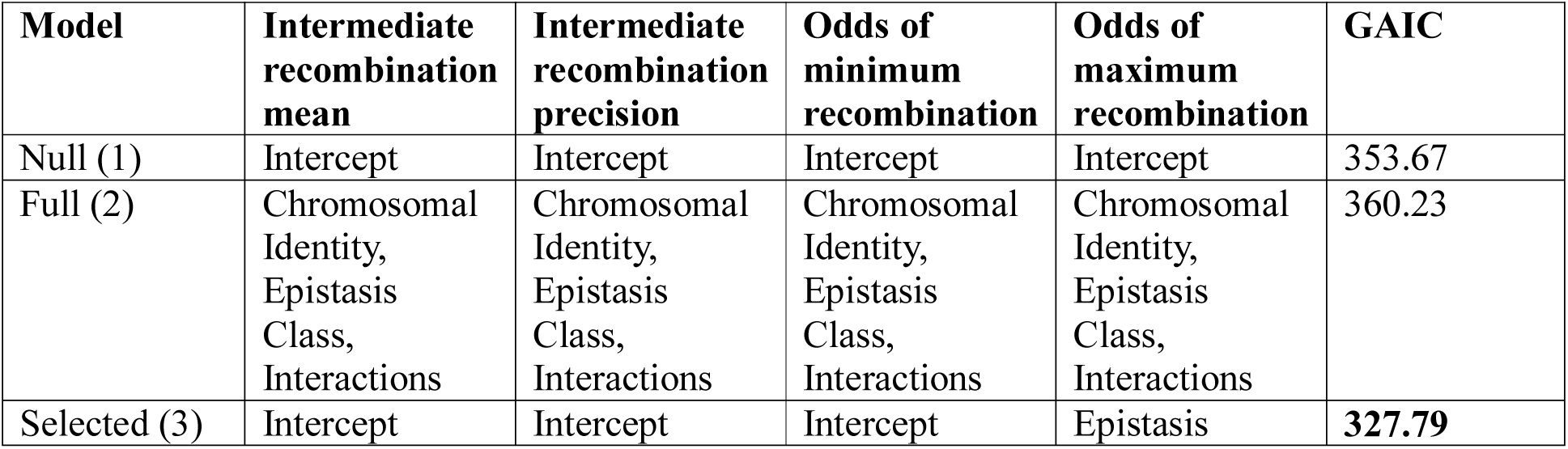
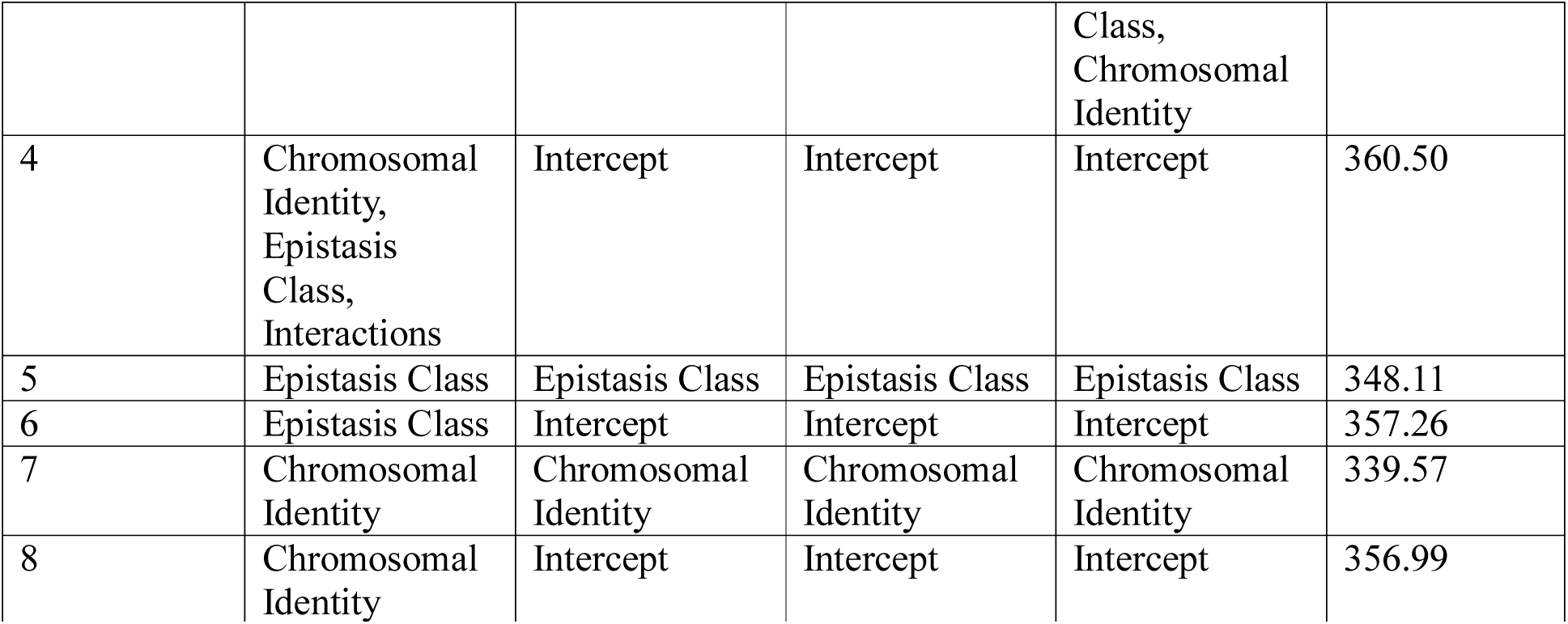
Comparison of models for the effect of chromosomal identity and epistasis class on the recombination rate between pairs of loci. Column headings refer to components defined in Methods (Flybase Meta-Analysis: Regression Model to Test for Effects of Epistasis Class and Chromosome Number on Recombination): 1) the mean of the intermediate recombination rates (i.e. recombination rates that were not 0 or 0.5); 2) the precision of the intermediate recombination rates 3) the relative probability that *c =* 0 rather than 0 < *c <* 0.5; and 4) the relative probability that *c =* 0.5 rather than 0 < *c* < 0.5. The names for them are: 1) Intermediate Recombination Mean, 2) Intermediate Recombination Precision, 3) Odds of Minimum Recombination, 4) Odds of Maximum Recombination. The final column shows the GAIC with the best fit (i.e. lowest value) in bold. The rows describe the variables included in each component of the model. An ‘Intercept’-only component means that none of the independent variables had an effect on that component of the recombination rate.

**TABLE 9:**
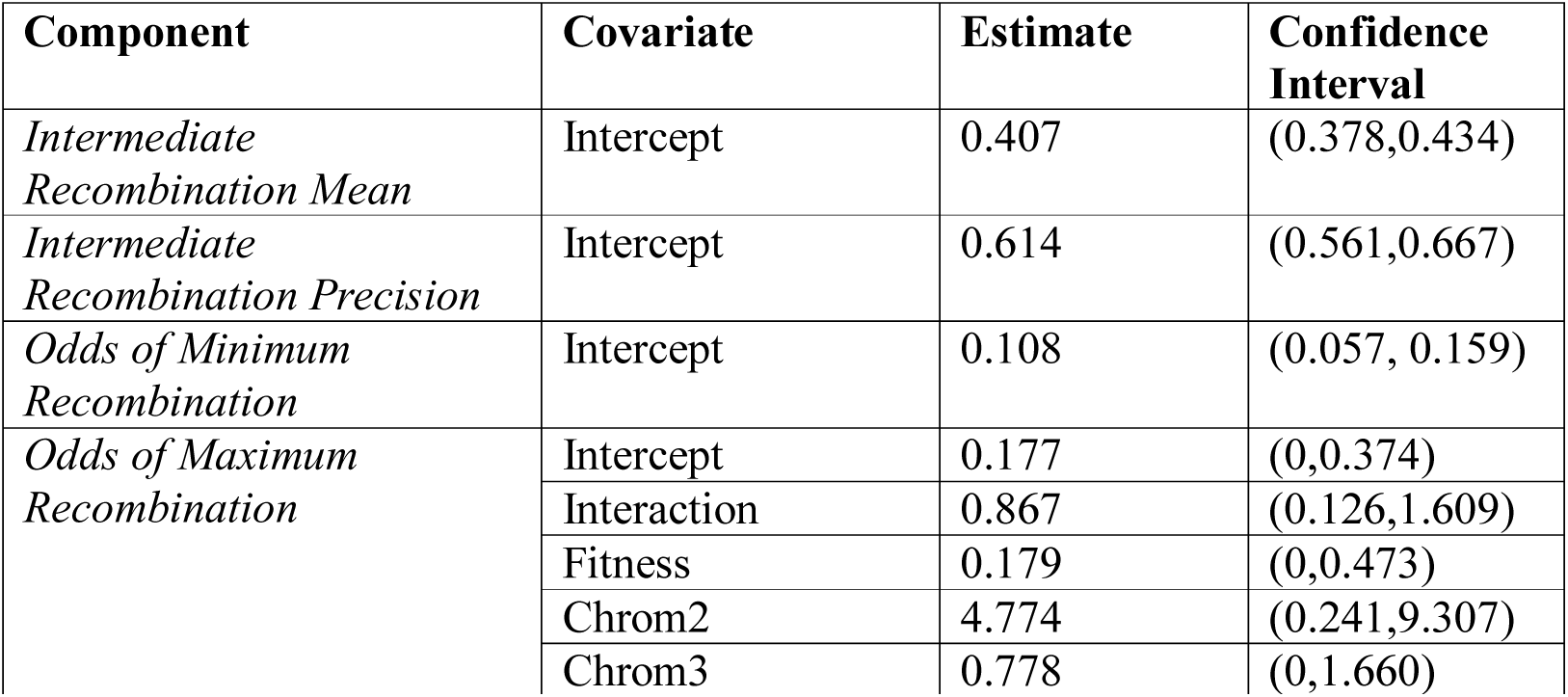
Best model for the effect of chromosomal identity and epistasis class on the recombination rate between pairs of loci. This is model 4 from [TABLE 8] & [FIGURE S2]. Column headings are variables included in the model, its parameter estimate (as defined in appendix II) and its standard error and 95% confidence intervals. Note that ‘component’ refers to any of the four parts of the response variable (recombination rate); these components are shown in the row headings and are defined in the Methods (Flybase Meta-Analysis: Regression Model to Test for Effects of Epistasis Class and Chromosome Number on Recombination): Intermediate Recombination Mean, Intermediate Recombination Precision, Odds of Minimum Recombination, Odds of Maximum Recombination. In contrast to ‘components’, ‘variable’ refers to any of the explanatory variables, that is the Intercept, Interaction, Fitness, Chrom2 and Chrom3 variables. The estimates given are transformed back to their original scale using the inverse of the link function and confidence intervals are calculated using the Delta Method.

| Component | Covariate | Estimate | Confidence Interval |
| --- | --- | --- | --- |
| *Intermediate Recombination Mean* | Intercept | 0.407 | (0.378,0.434) |
| *Intermediate Recombination Precision* | Intercept | 0.614 | (0.561,0.667) |
| *Odds of Minimum Recombination* | Intercept | 0.108 | (0.057, 0.159) |
| *Odds of Maximum Recombination* | Intercept | 0.177 | (0,0.374) |
|  | Interaction | 0.867 | (0.126,1.609) |
|  | Fitness | 0.179 | (0,0.473) |
|  | Chrom2 | 4.774 | (0.241,9.307) |
|  | Chrom3 | 0.778 | (0,1.660) |

CAPTION: [TABLE 9] : Best model for the effect of chromosomal identity and epistasis class on the recombination rate between pairs of loci. This is model 4 from [TABLE 8] & [FIGURE S2]. Column headings are variables included in the model, its parameter estimate (as defined in

The best fitting model (number 4 in [TABLE 8]) appeared to be a good fit (GAIC = 327.79); each component sub-model except the Odds of Maximum Recombination component only included the Intercept, and the Odds of Maximum Recombination component included the Fitness, Interaction, Chrom2 and Chrom3 variables but no interaction variables. In other words, a constant model was sufficient to explain the distribution of recombination rates of locus pairs except for locus pairs which recombined at the maximum rate. Within the Odds of Maximum Recombination component, only the Fitness variable was significant (transformed effect: 0.179, 95% CI: (0,0.473)). We considered an effect significant if it appeared in both the transformed scale and the original scale, i.e. it was robust to being transformed as discussed in Methods.

### Simulations

In the simulations performed, the fixation probability of the beneficial genotype could display very different behaviours depending on the recombination rate, how close the initial population was to linkage equilibrium, and how many individuals bore the beneficial haplotype. [FIGURE 10] gives a graphical description of the behaviour with different initial population distributions, all with a population size of 200. In all cases the beneficial haplotype was (0,0). In each raster the x-axis is the recombination rate used for each simulation, the y-axis is the 1 + s in each simulation, and the colour is the probability of fixation of (0,0). The table in the right-hand corner gives the initial number of each haplotype in every simulation depicted. Higher rates of selection always increased the probability of the fixation of the beneficial haplotype, which is an intuitive and well-known result. When the population started close to linkage equilibrium [FIGURE 10, Case 2], recombination had no effect on the probability of fixation. When the population started far from linkage equilibrium, and had a small number of individuals of the beneficial genotype *and* a relatively small number of individuals bearing the non-beneficial coupling haplotype (1,1), higher rates of recombination gave a higher probability of fixation [FIGURE 10, Cases 1 & 10]. When the initial population started far from linkage equilibrium, and had at least one individual bearing the beneficial haplotype and a large number of individuals bearing the non-beneficial coupling haplotype, higher recombination rates reduced the probability of fixation [FIGURE 10, Case 5 and 12]. These patterns were also reflected in the simulations with large recombination rates [FIGURE S3].

## Discussion

The results of the meta-analysis indicate that the odds of a locus pair belonging to the PDFI epistasis class with a deleterious fitness effect recombining at the maximum rate over recombining at an intermediate rate are is about one-fifth the corresponding odds of a pair belonging to the PDNF class (odds ratio: 0.179, 95% CI: 0 – 0.473, see [TABLE 9]). The confidence interval of this effect is very large, which is most likely due to a relatively small number of PDFI loci (36), but nonetheless seems to be strong evidence of an effect.

Two possible explanations for the reduced recombination rate are as follows. First, the non-lethal interactions are more likely to be maintained in regions of low recombination, possibly because if they occur in a region of high recombination, they would be quickly broken up. Second, regions with this kind of deleterious interaction experience selection against recombination, resulting in reduced recombination in that region. The two explanations are not mutually exclusive.

We did not observe a difference between epistasis classes when comparing locus pairs that did not recombine at all (*c = 0*) to pairs that recombined in the range (0,0.5) [TABLE 9]. A possible explanation for this is that there is no qualitative difference between zero recombination and an intermediate recombination rate when it comes to lethal interactions – all that matters is whether a locus pair recombines at the maximum rate or not. Another possibility is that we simply did not detect a difference where it existed, due to the relatively small number of locus pairs with zero recombination rate vs those recombining freely (19 vs 51).

There were a number of non-significant variables in the model which we included out of interest: the Chrom2 variable increased the odds of a locus pair recombining at the maximum rate over recombining at an intermediate rate (transformed value: 4.774); on the other hand, the Chrom3 and Interaction variables all decreased these odds (with transformed values of 0.778, 0.867 respectively).

It is important to note in the studies we curated and analysed, the selection pressures have to be quite strong to be detected. There is some evidence from mathematical modelling that other weaker epistatic fitness effects could show different patterns but such effects are much more difficult to detect than lethality.

Our simulations indicate that there is a broad range of cases in which a high recombination rate is advantageous and another broad range of cases in which it is not. This is consistent with the lack of consensus in the previous literature. Whether recombination is advantageous or not is mainly controlled by whether there is a substantial number of the beneficial genotype in the initial population. If initially there is a large number of beneficial genotypes, recombination is less advantageous [Figure 10, “Case 3” & “Case 6”]. The latter is also determined by the relative proportion of repulsion genotypes. If repulsion genotypes are common recombination becomes advantageous [Figure 10, “Case 9”, “Case 11” & “Case 12”]. This means that the threshold between advantageous and disadvantageous occurs at linkage equilibrium, which agrees with prior mathematical literature (Mano 2013).

**FIGURE 10:**
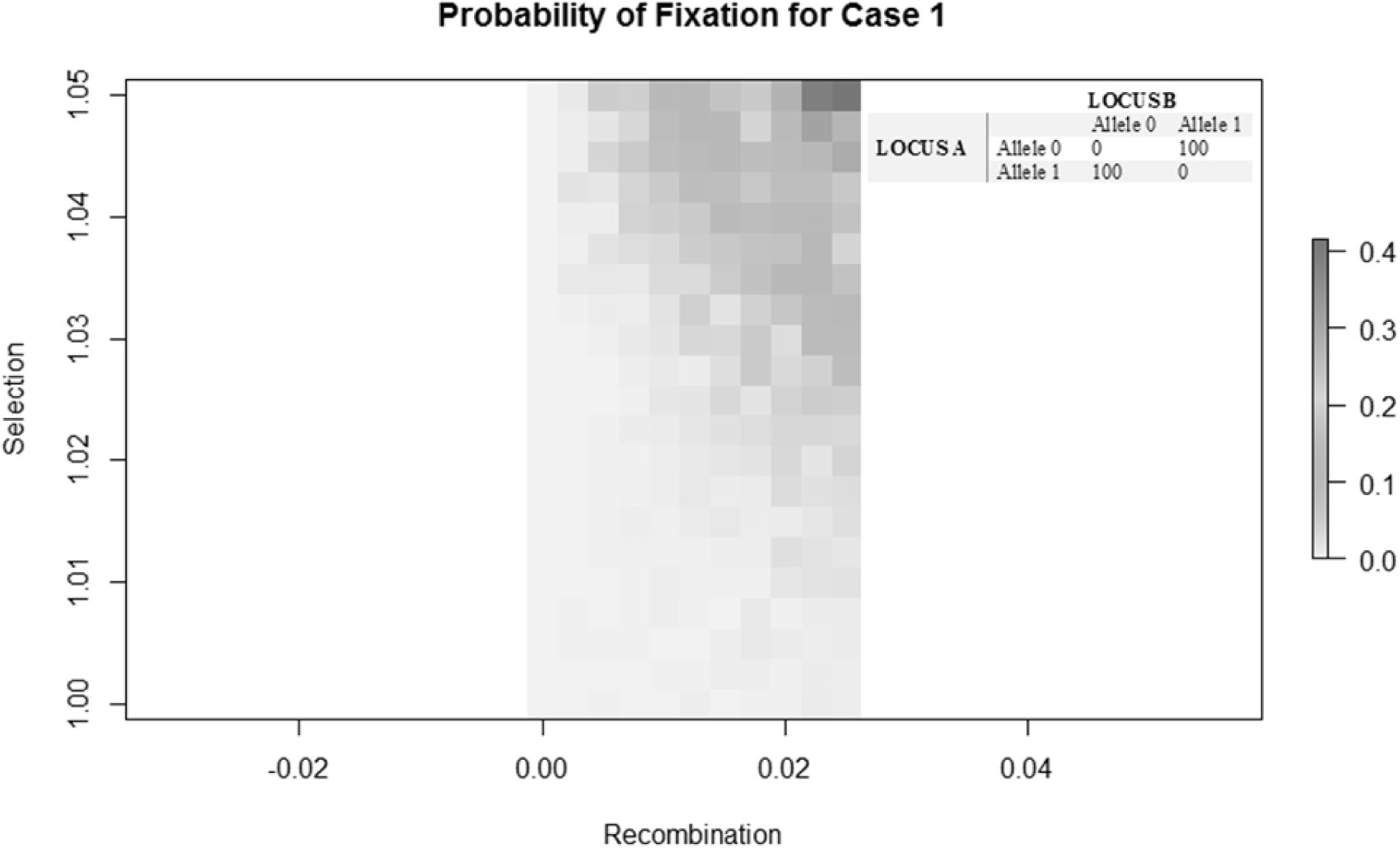
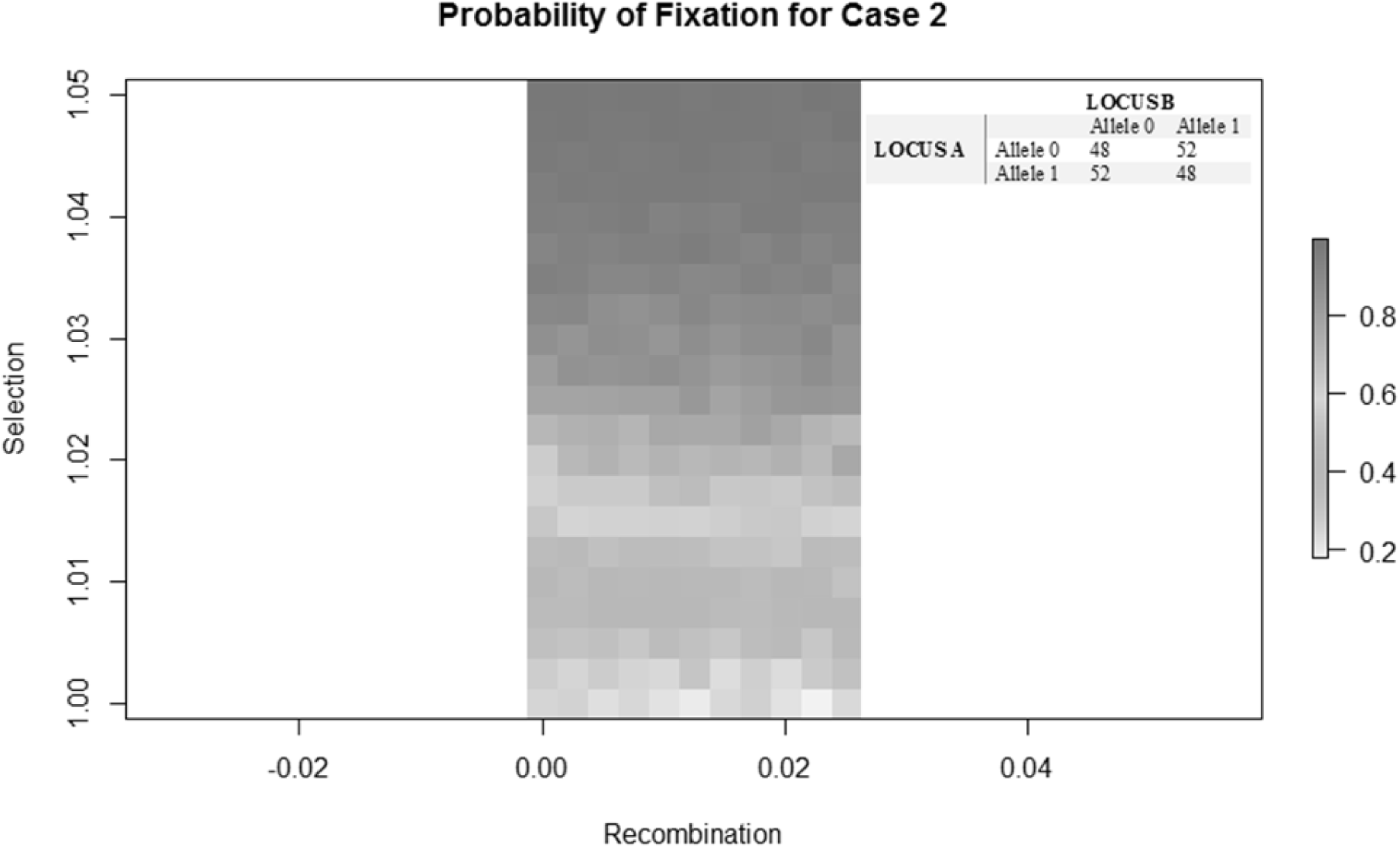
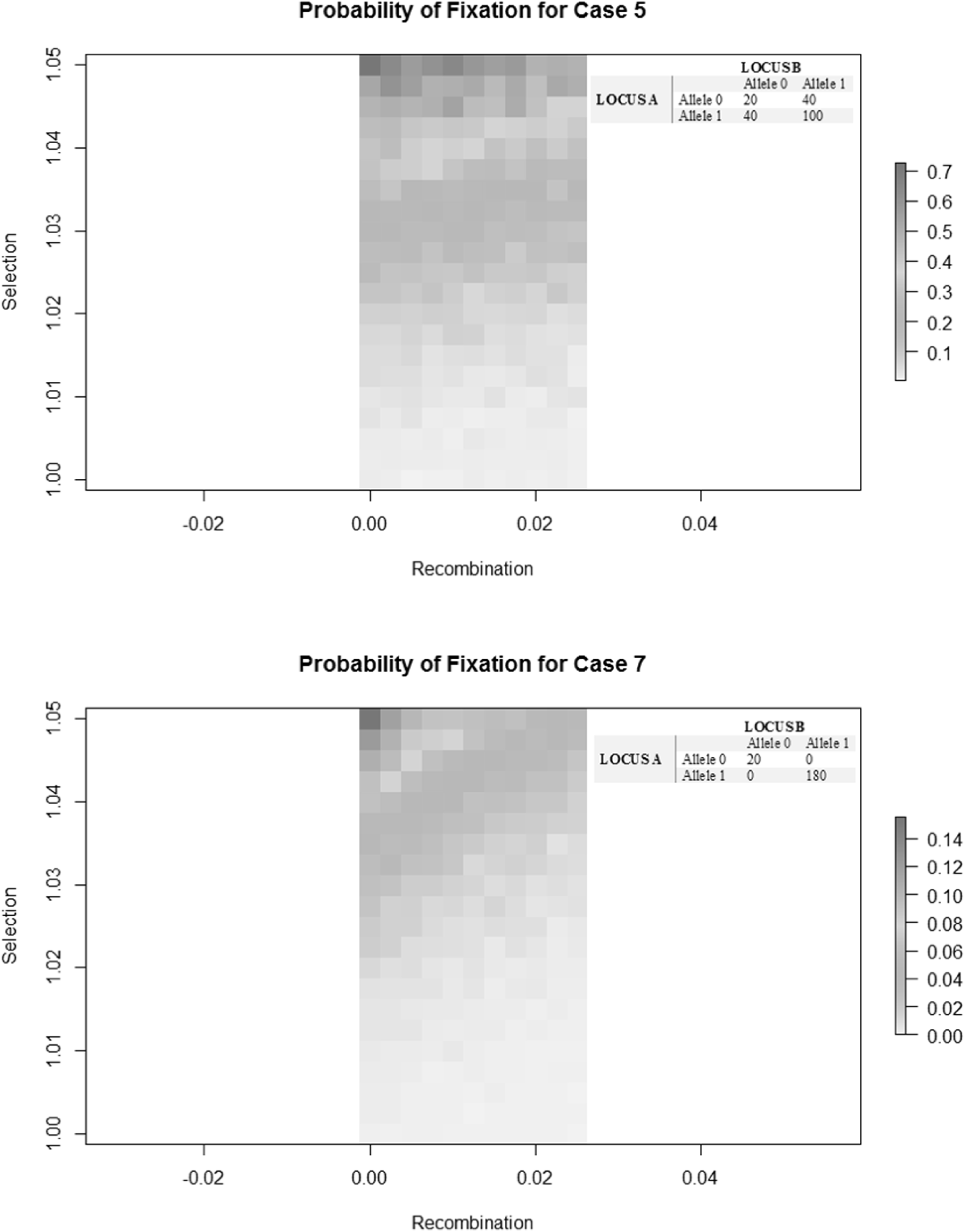
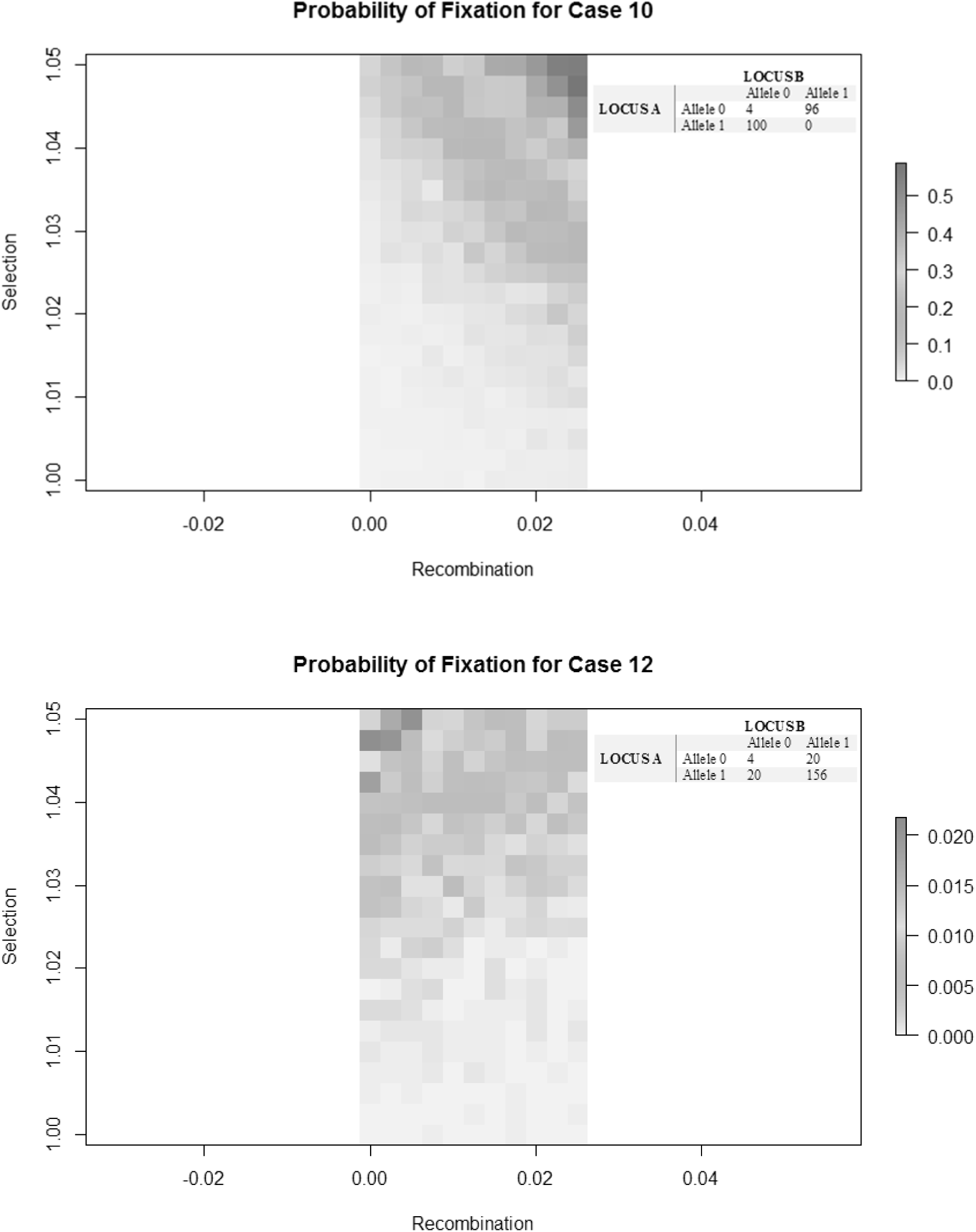
Probability of fixation on two axes – recombination and selection. Each column of shading is centred on a recombination value on the horizontal axis. In all cases the beneficial haplotype was (0, 0). A darker colour indicates a higher probability of fixation (and thus advantage of that recombination and selection combination). The table in the top right corner gives the initial population counts. All graphs are of populations with 200 individuals. The Supplement gives more plots for the results of the cases not shown here [FIGURE S3].

In our simulations the effect of beneficial epistatic interactions could favour increased rates of recombination in cases where the population was far from linkage equilibrium and there were not too many (relative to population size) of either the beneficial haplotype or the non-beneficial coupling haplotype. The effect of linkage disequilibrium is likely explained by the fact that, in the neutral case, the marginal distributions of the alleles evolve independently – that is, a population at linkage equilibrium is likely to stay in linkage equilibrium even as it moves to fixation in one of the types (Durrett and Mano 2013). Our simulations suggest that this remains (roughly) true even with epistatic selection. This is because for those of our simulations where the population was initially in linkage equilibrium, the fixation probabilities at each locus decoupled and became independent of one another at 50 generations, as would be expected of a population that remains in linkage equilibrium. This can clearly be seen in the fixation probability graphs of these cases, where there are horizontal rather than diagonal lines visible, indicating that fixation probability no longer depends on recombination rate. [Figure 10, “Case 2”, “Case 3”, “Case 4” & “Case 5”]

The reduced effectiveness of recombination in facilitating selection when there are several of the non-beneficial coupling haplotype is most likely because the beneficial haplotype and the non-beneficial coupling haplotype recombine to make the repulsion haplotypes, meaning that on average there will be fewer individuals bearing the beneficial haplotype than if there were no recombination.

Results from our simulations and meta-analysis were complementary, as genotypes from the meta-analysis were lethal but beneficial genotypes were studied in the simulations. There was broad agreement between the two - *D. melanogaster* genotypes with lethal phenotypes displayed tight linkage, which is very similar what our simulations indicate would occur in a population with a large number of a beneficial genotype ((0,0)) or the genotype which opposes it ((1,1)) [FIGURE 10, “Case 7” & “Case 8]. In both cases non-beneficial or lethal genotypes very quickly reduce in frequency and are eliminated from the population with little to no recombination.

It is not clear how well the meta-analysis results will generalise to other species. Less closely related organisms are likely to display different recombination and selection patterns to *Drosophila melanogaster* (Dumont and Payseur 2008). In particular, mammals have so-called ‘recombination hotspots’ throughout the genome, with regions of size 1-2 kilobases displaying very high rates of recombination, but with very large regions with very little recombination. In humans the recombination rate within such a hotspot can be as high as 50 times the average of the rate over the entire chromosome (Kauppi *et al*. 2004). *Drosophila melanogaster* does not display such punctuated recombination rates (Comeron *et al*. 2012).

Population size reductions are a common occurrence, happening regularly as part of a cyclical process, or in response to some kind of threat such as habitat loss. A population would be more likely to persist if beneficial combinations of alleles lost to such a population size reduction were regenerated via recombination within a few generations. This would require relatively high rates of recombination between pairs of loci that interact to have an effect on fitness. Unfortunately, our meta-analysis of *D. melanogaster* data suggests that the opposite is in fact the case – that in fact the rates of recombination between pairs of loci with a fitness interaction are particularly low [RESULTS TABLE]. The simulations show conditions under which recombination would be an advantage, which depend upon linkage disequilibrium and allele proportions for loci with epistatic effects on fitness. As more of these loci are discovered in natural populations of non-model species, it will be useful to assess whether these species are likely to experience effects that favour or select against recombination between epistatically interacting loci. This will have important implications for evolution in general, and for conservation management, because, as stated above, population persistence would be more likely if beneficial combinations of alleles lost during a population size reduction were regenerated rapidly via recombination.

This study has taken some of the first steps in empirically analysing the relationship between epistatic selection and recombination, but much more needs to be done, because epistatic selection and recombination are both ubiquitous and their interaction is both important and poorly understood.

Experimental work on this type of problem may be difficult, given how difficult it is to identify epistatic genetic interactions (Chanda *et al*. 2007), but it may be possible to do a similar analysis on other model species as more data becomes available. Studies of speciation may also yield some important results as experimental evidence has shown that hybrid male sterility is often controlled by epistatic interactions (Palopoli *et al*. 2004, Chang *et al*. 2010).

## Author Contributions

William Sherwin, John Murray, Mark Tanaka and Beniamin Goldys initiated this project as part of a UNSW mathematical biology initiative. Antony Bellanto carried out preliminary investigation of Flybase. Luis Cayetano performed preliminary simulations.

## Acknowledgements

Some early programming work was performed by Julian Marchal and Camilo Cassel.

## Code Supplement

### I. Input and Search Patterns

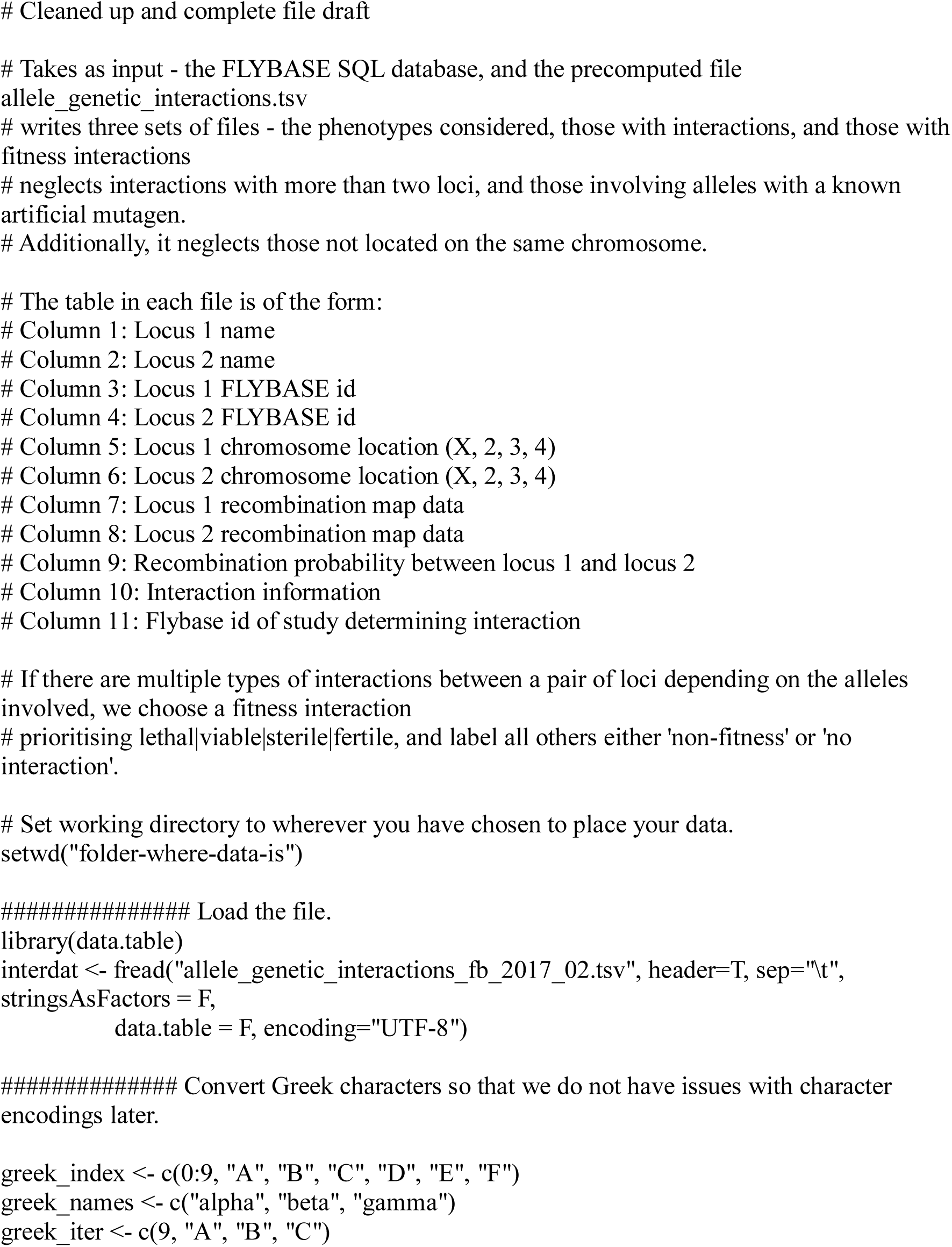
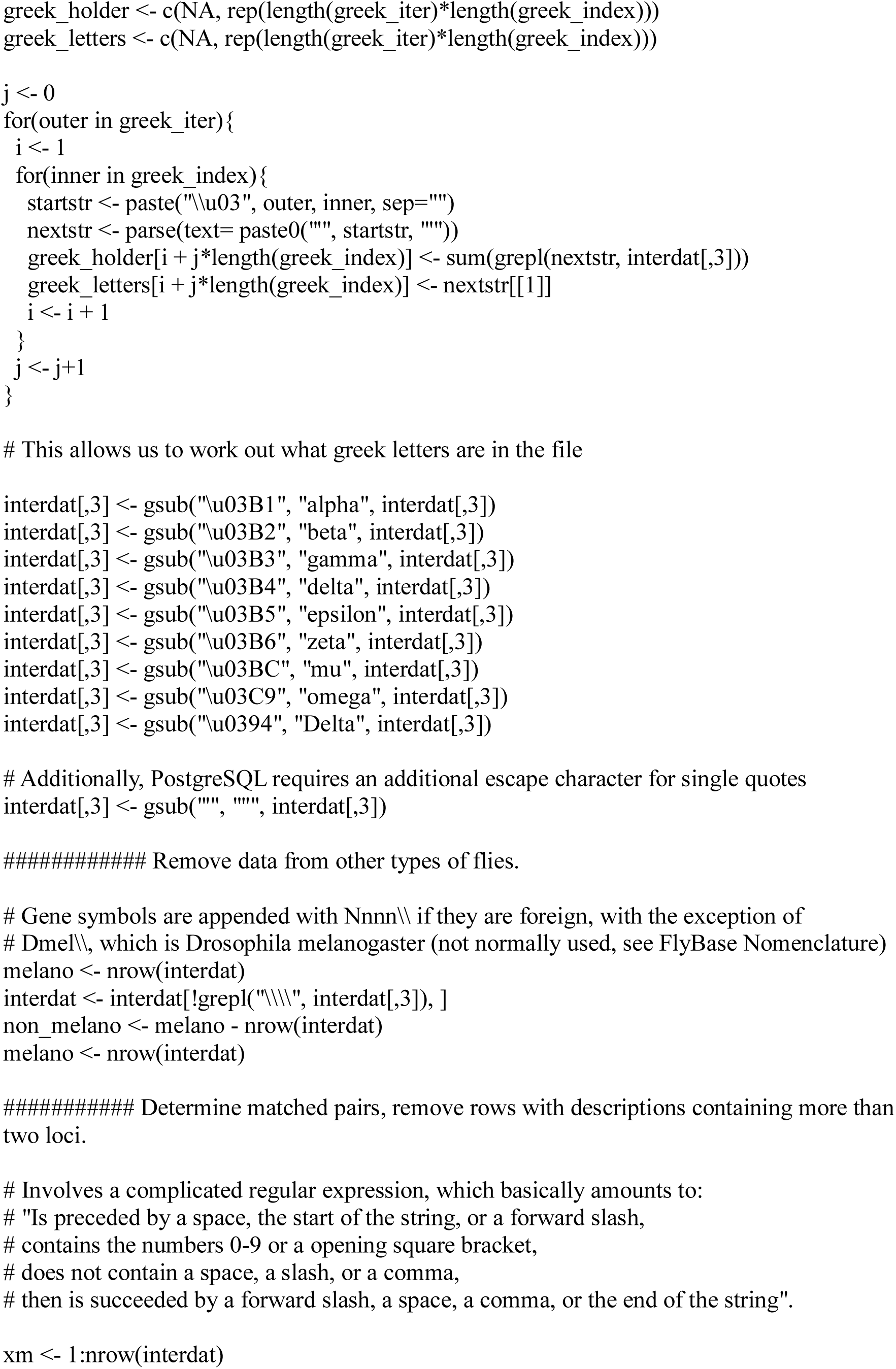
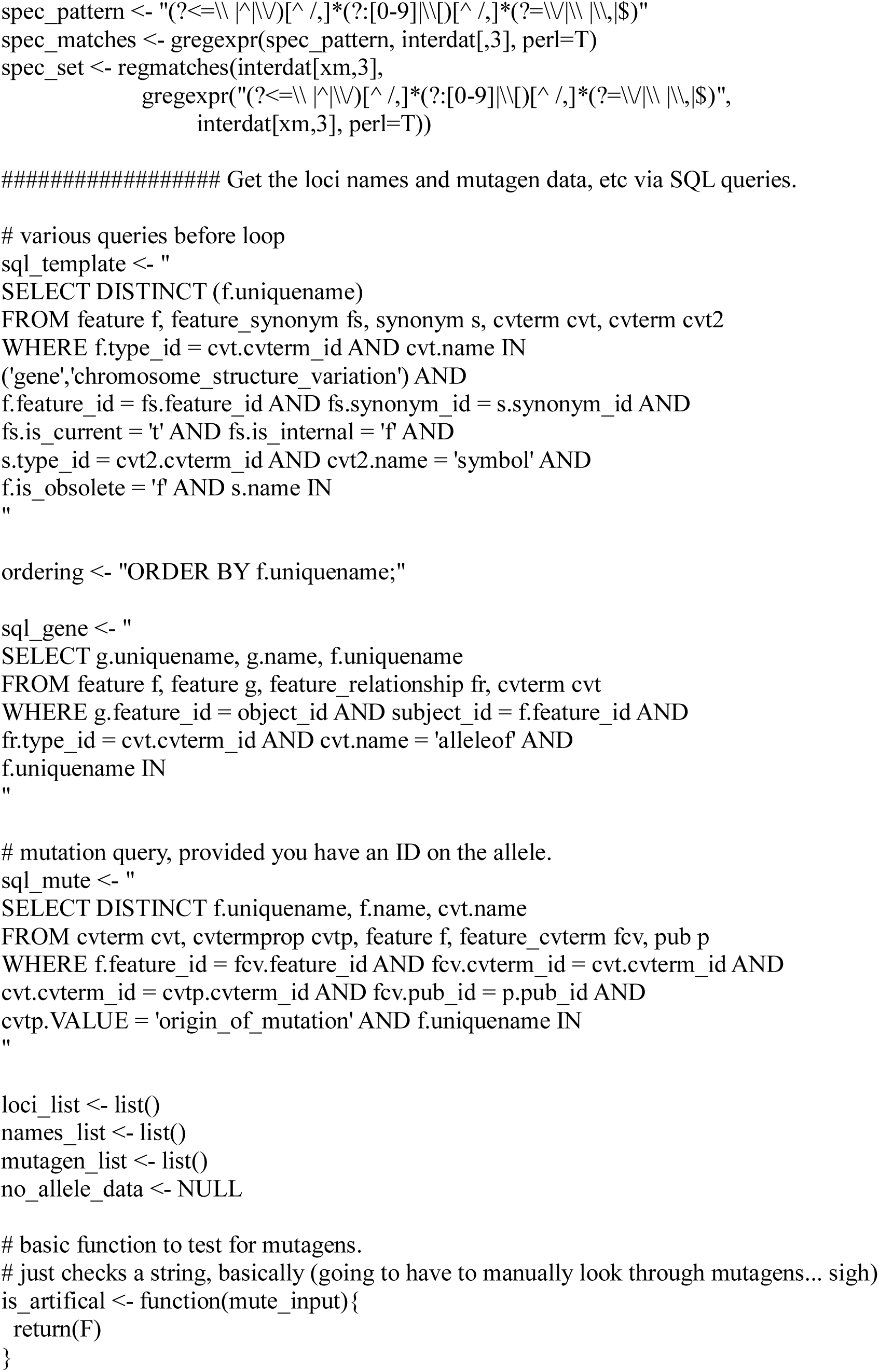
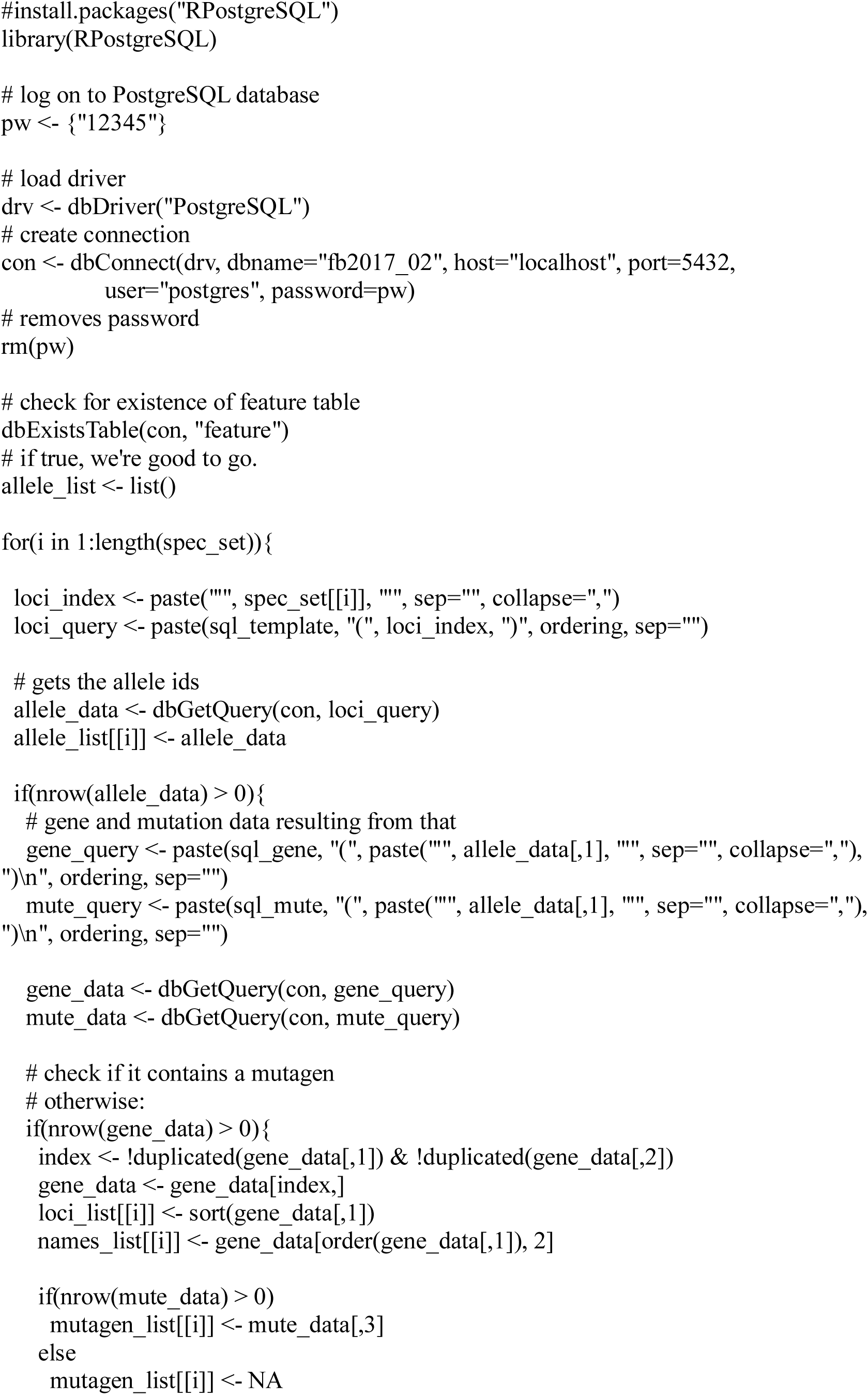
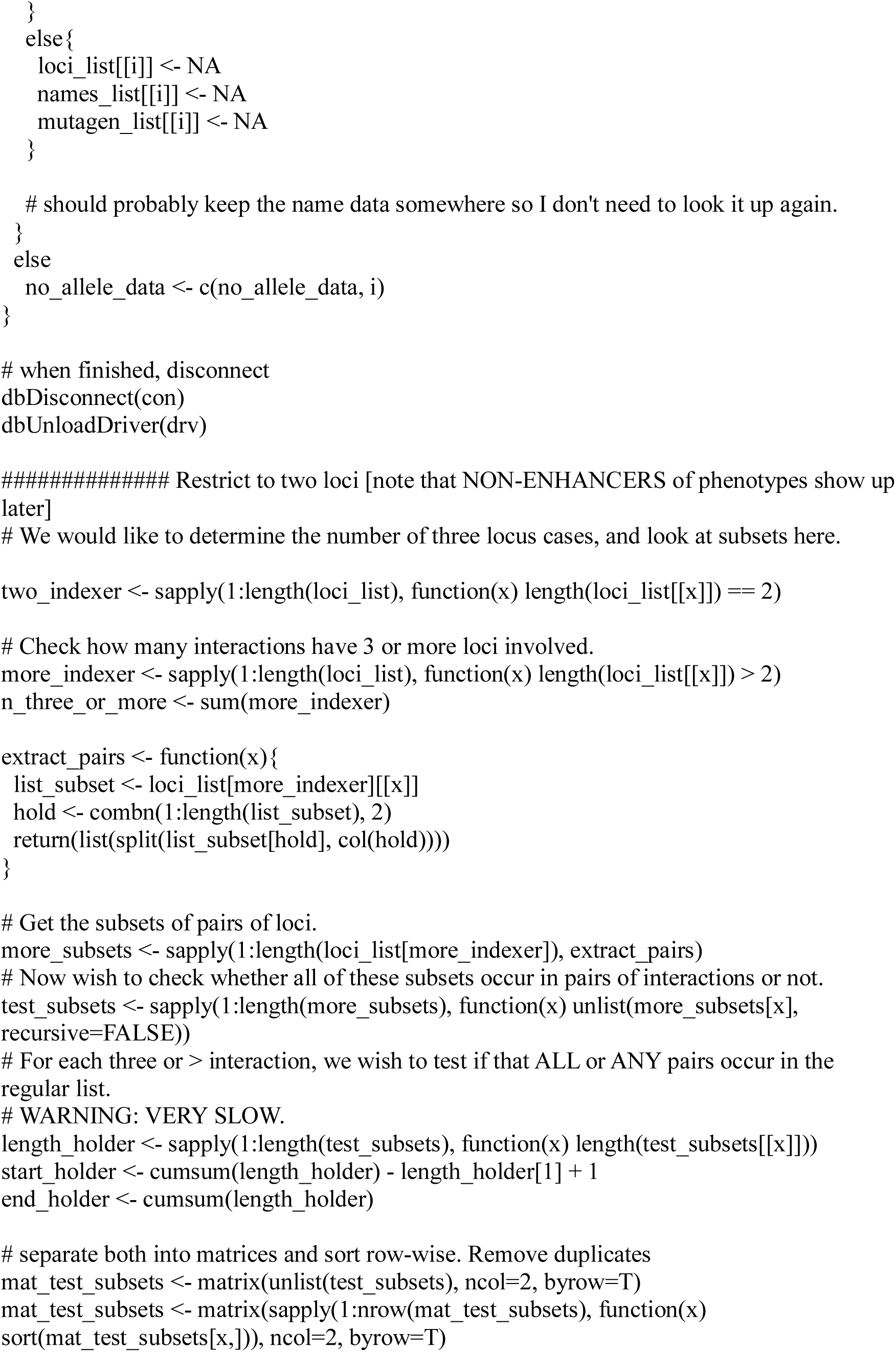
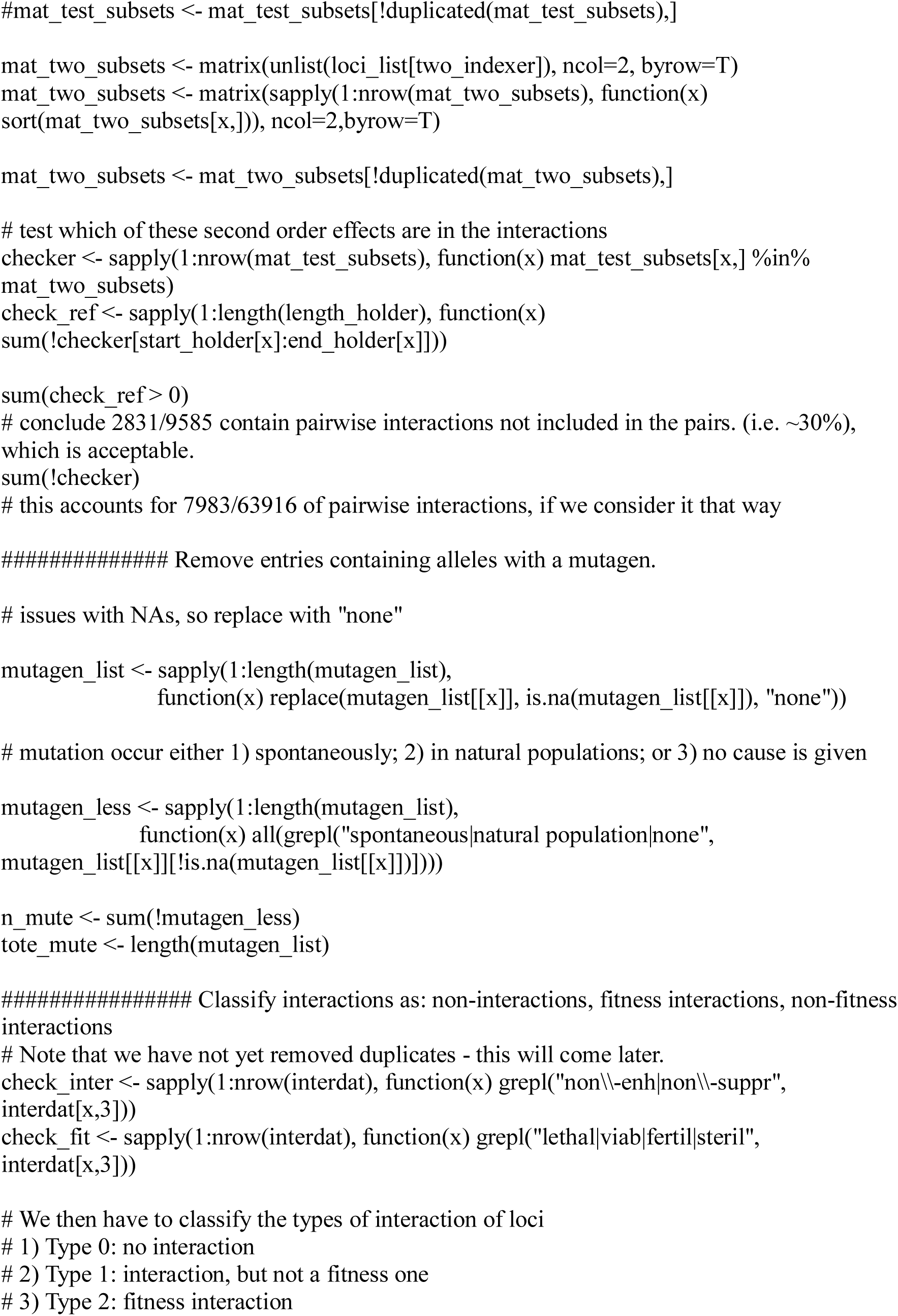
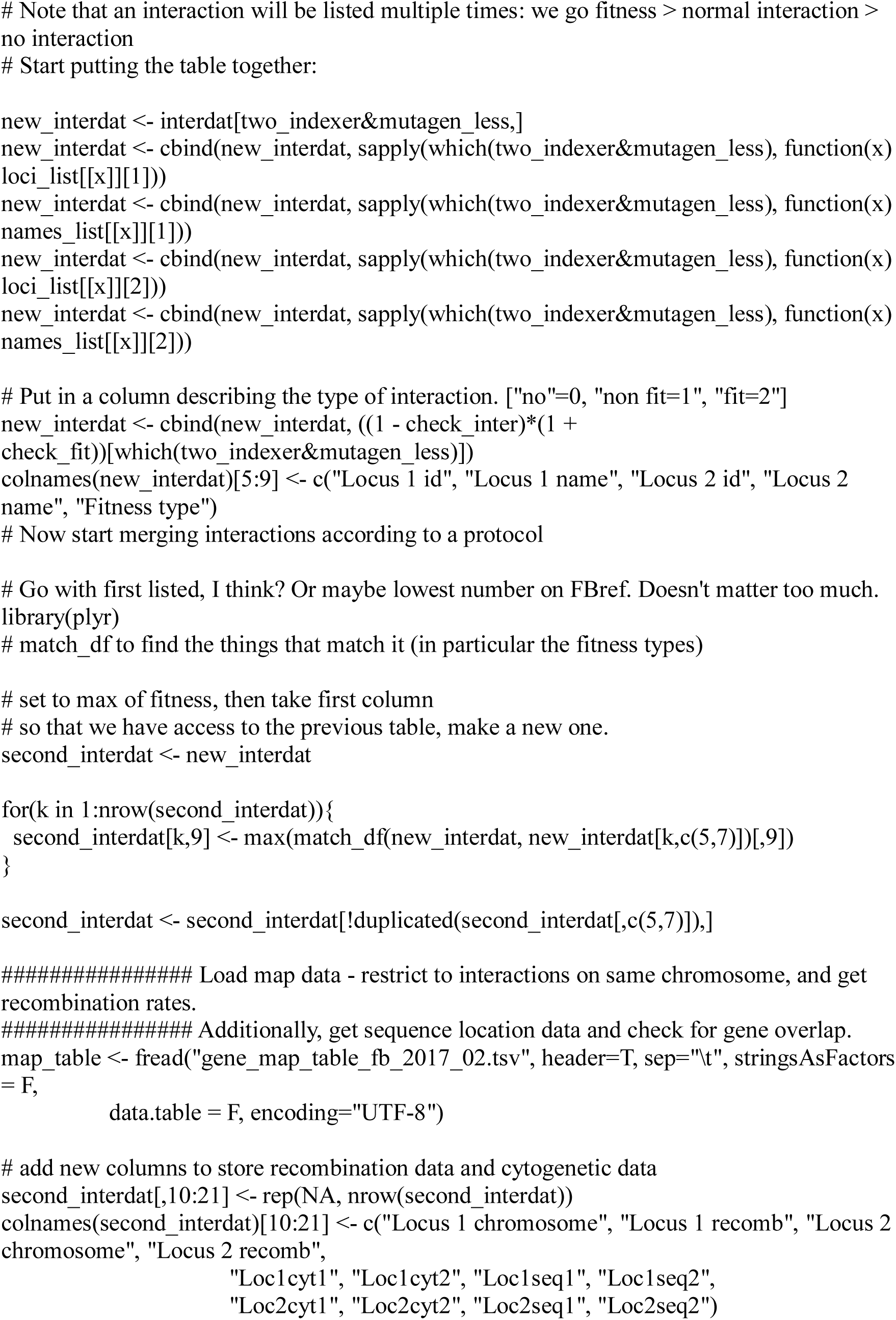
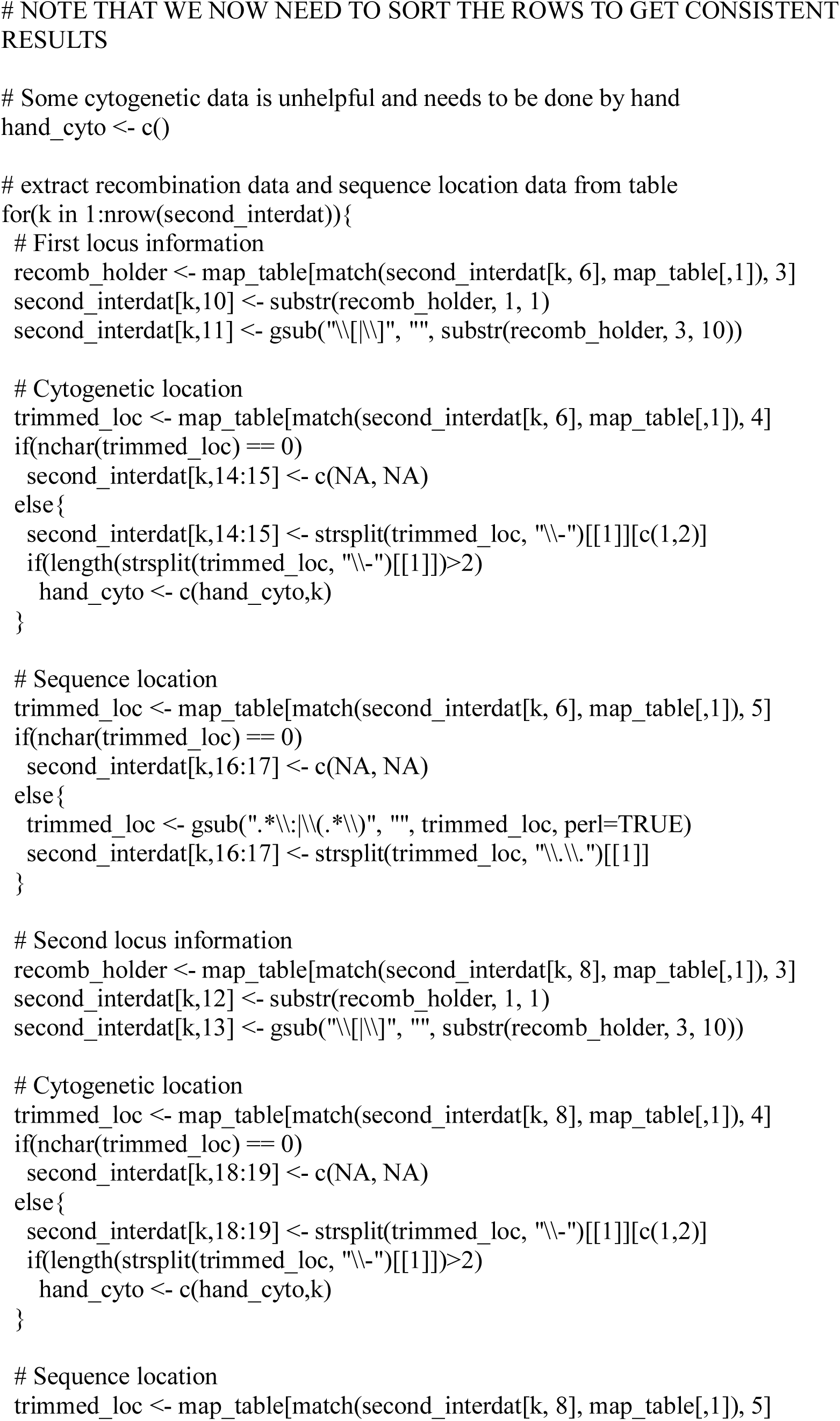
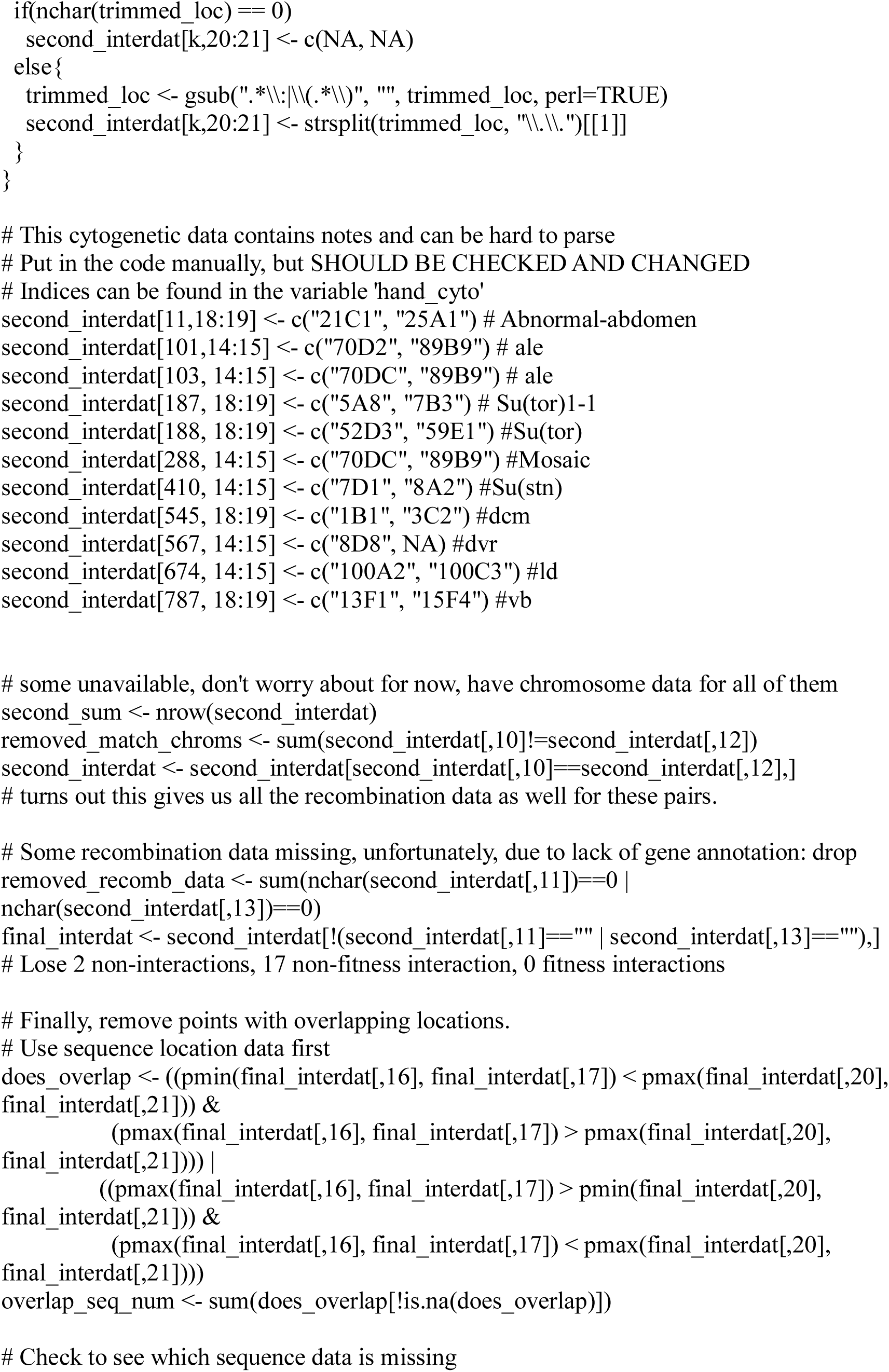
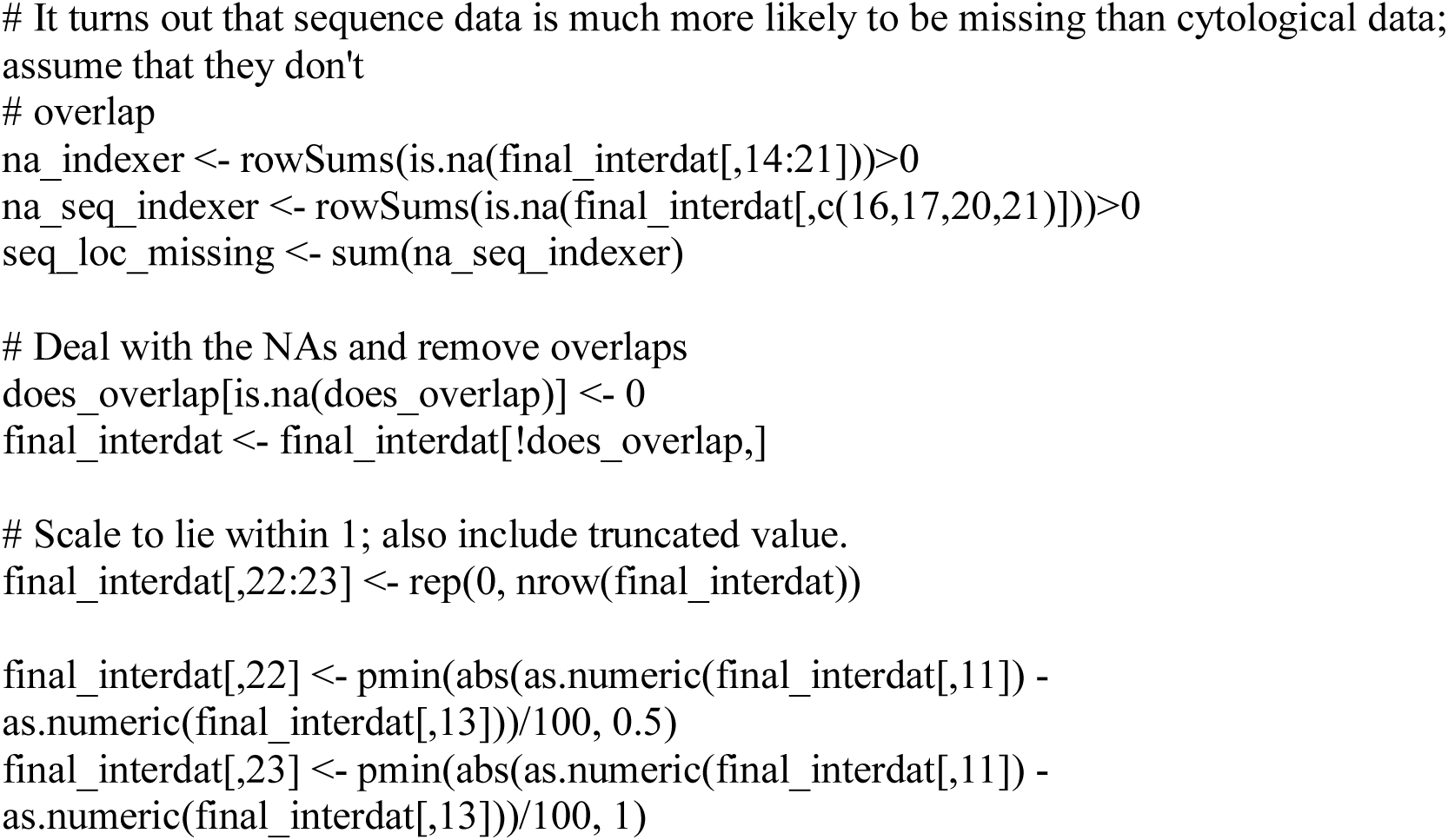

### II. Data Analysis

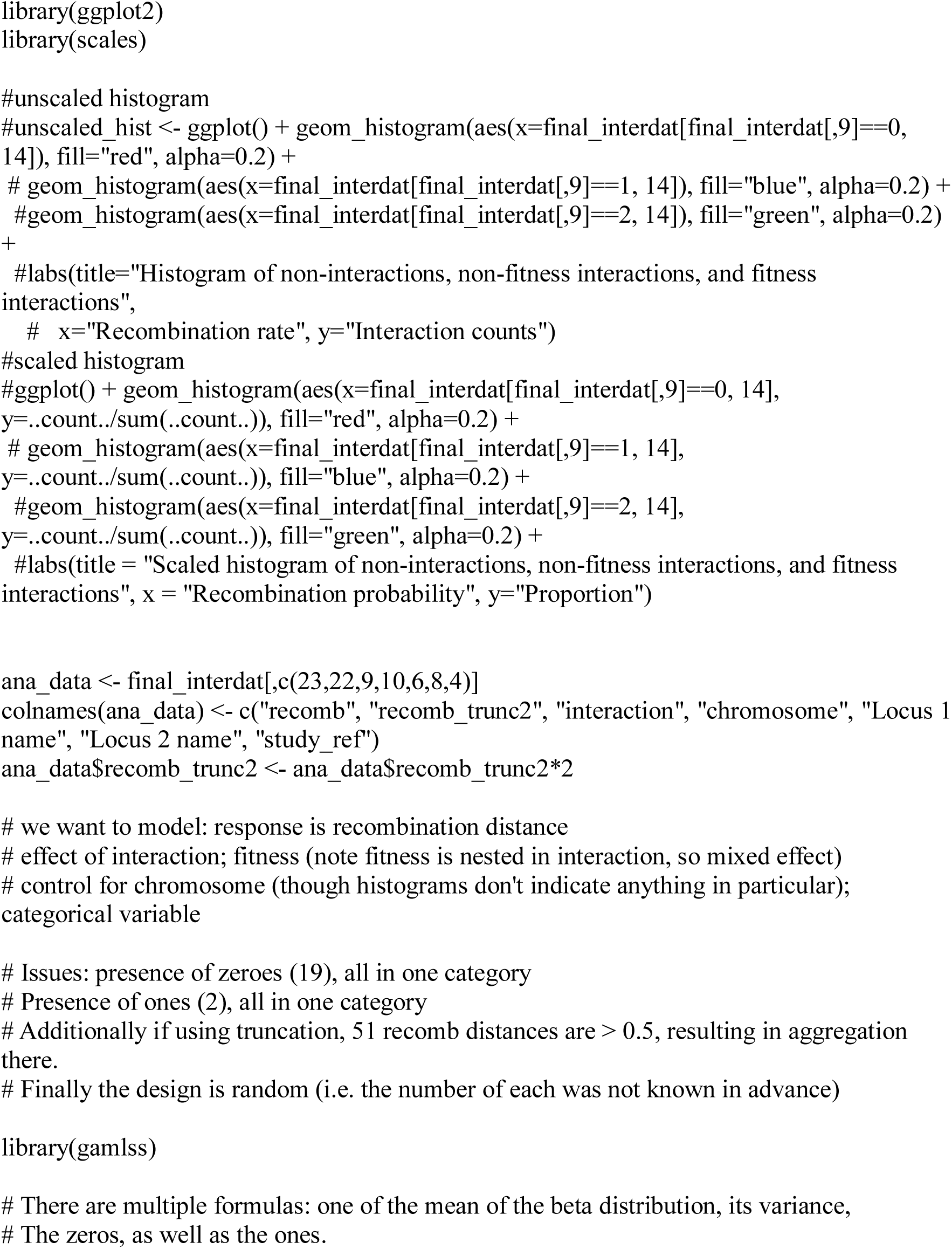
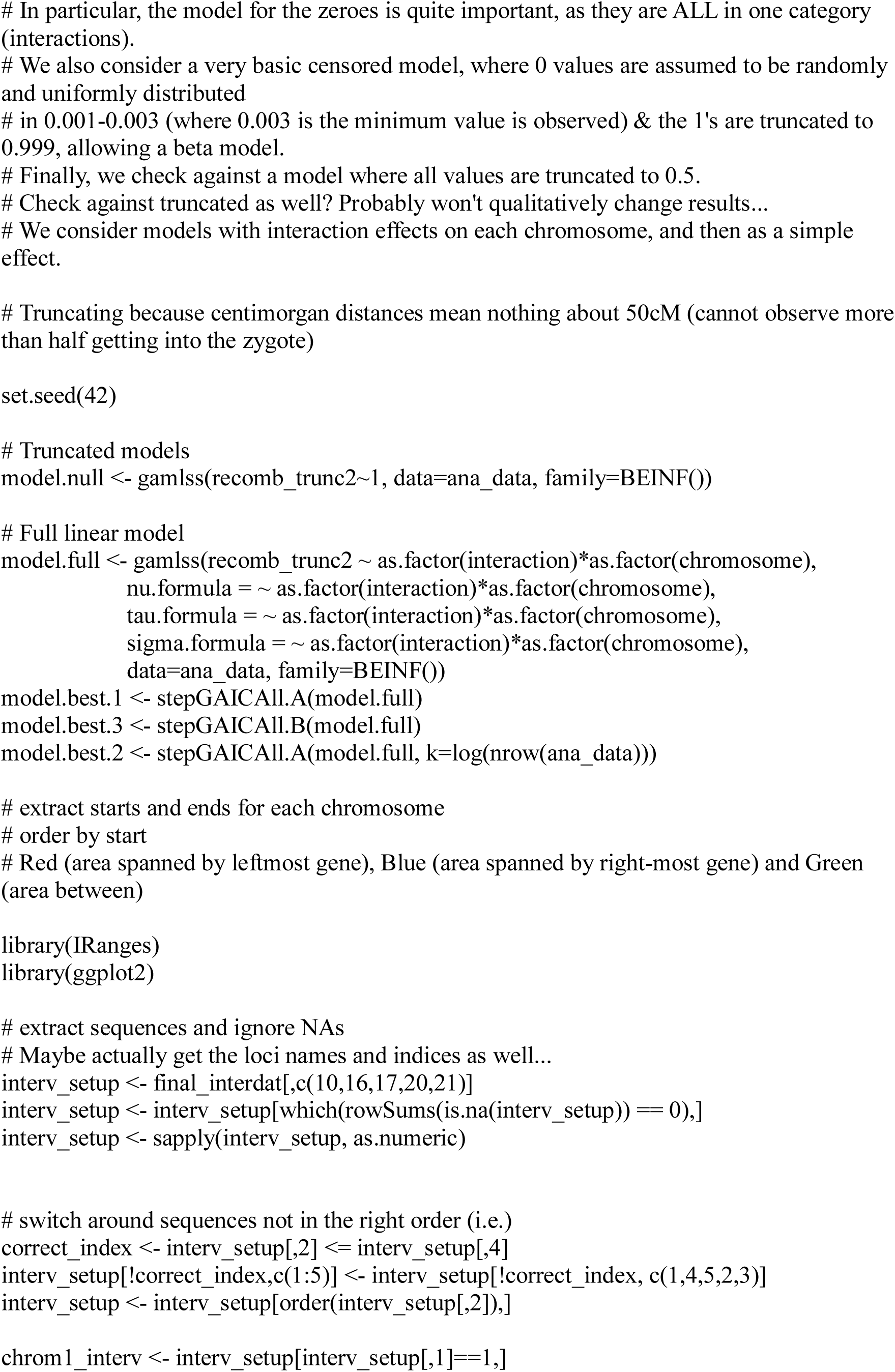
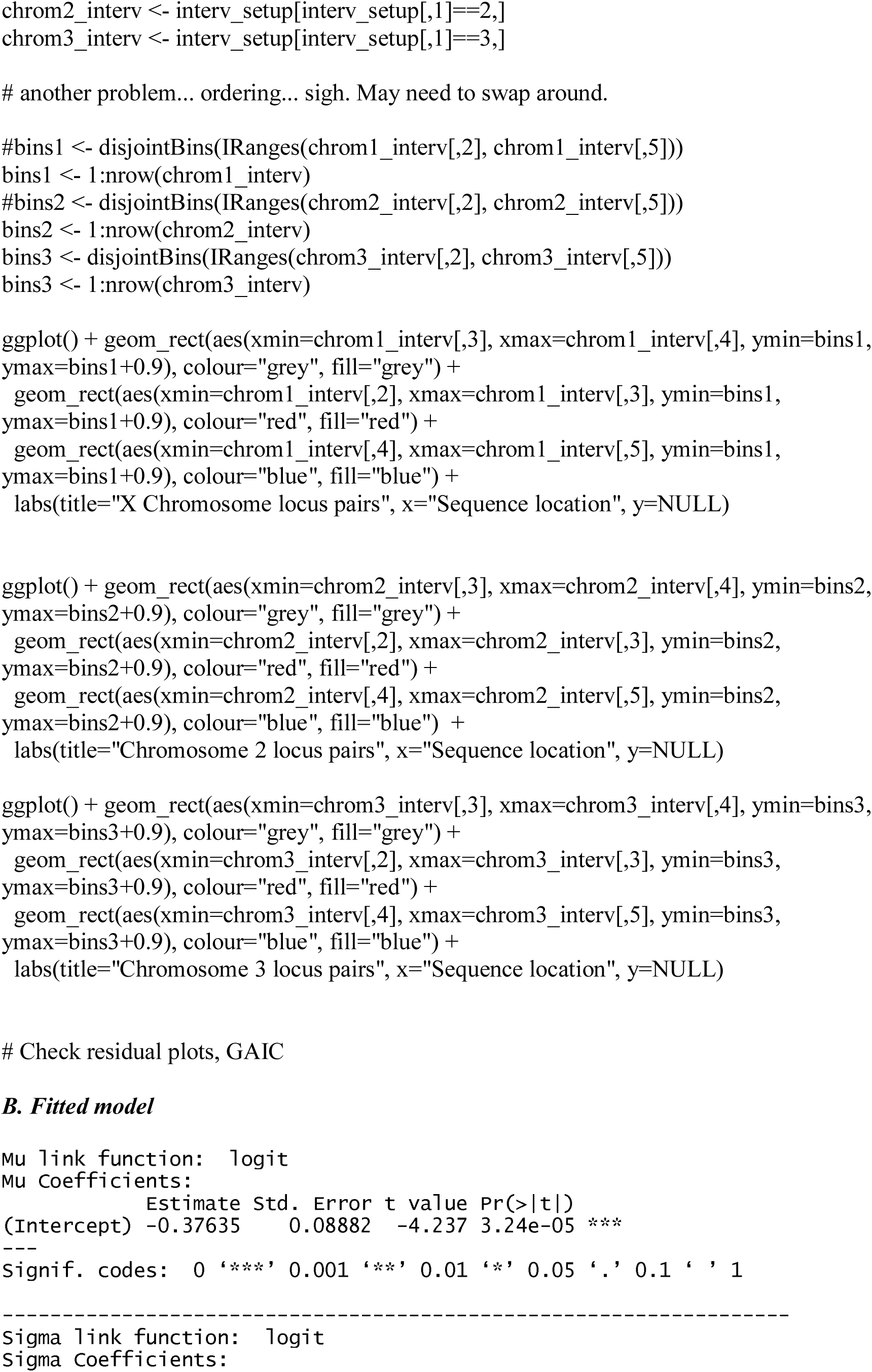
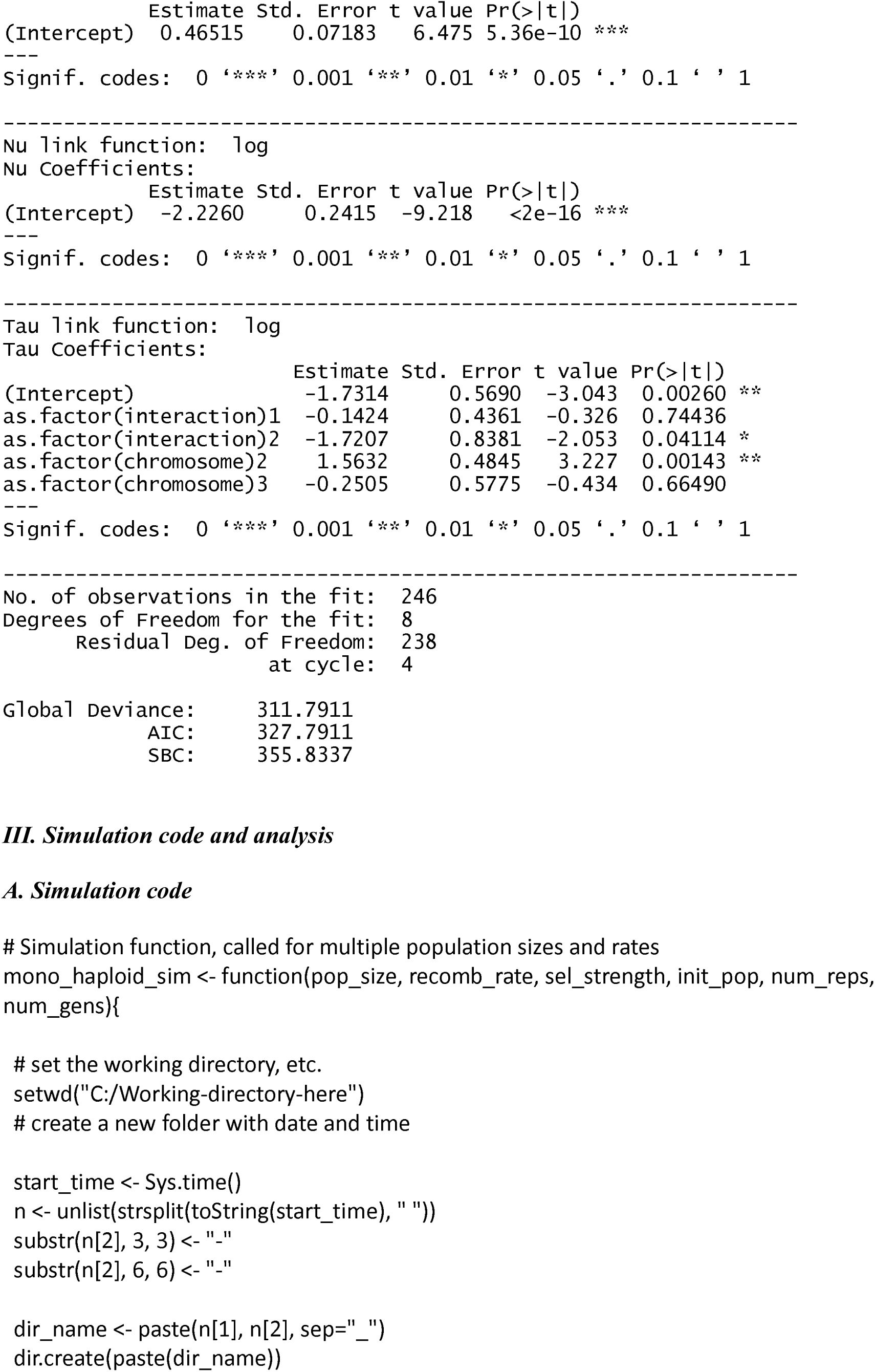
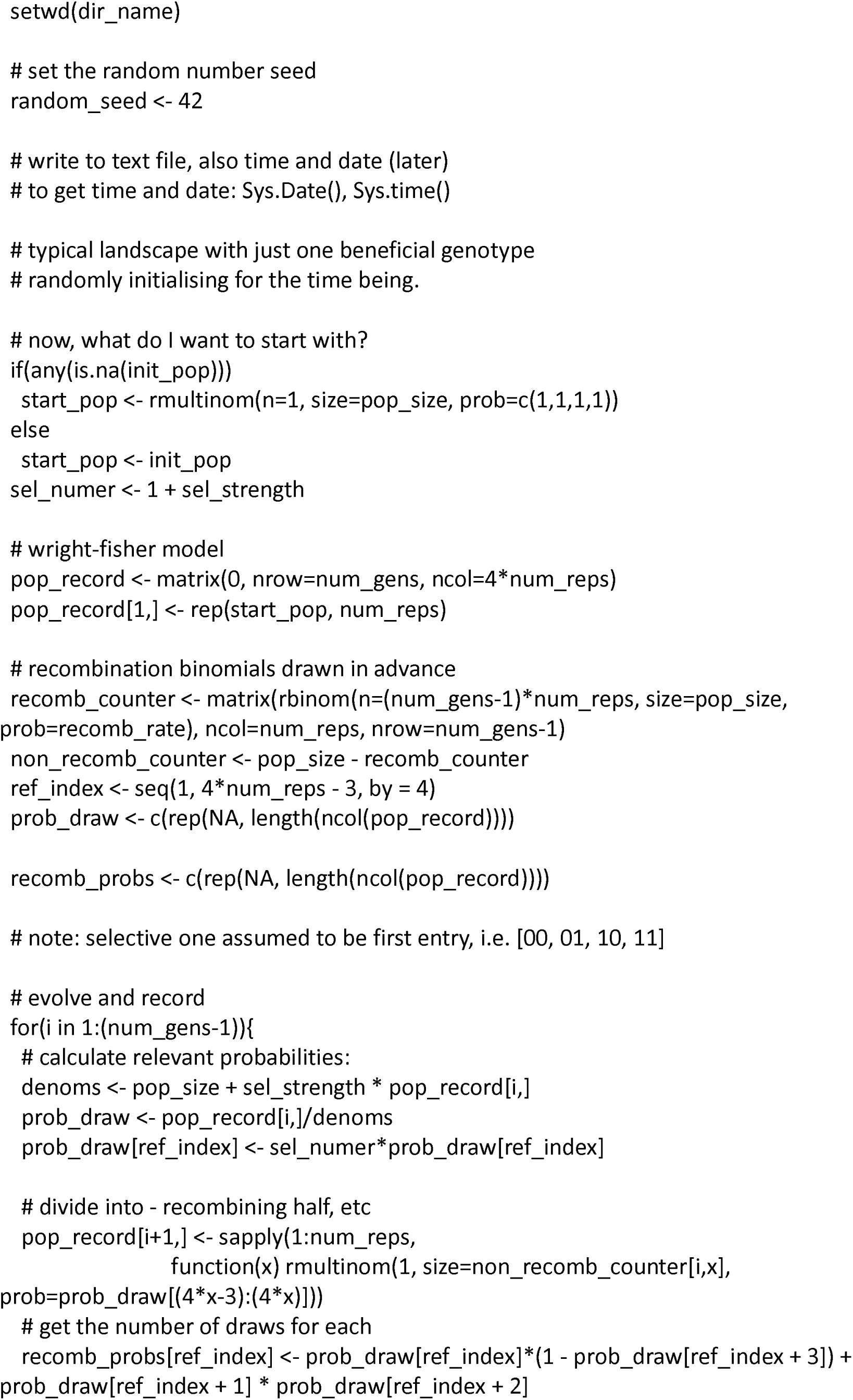
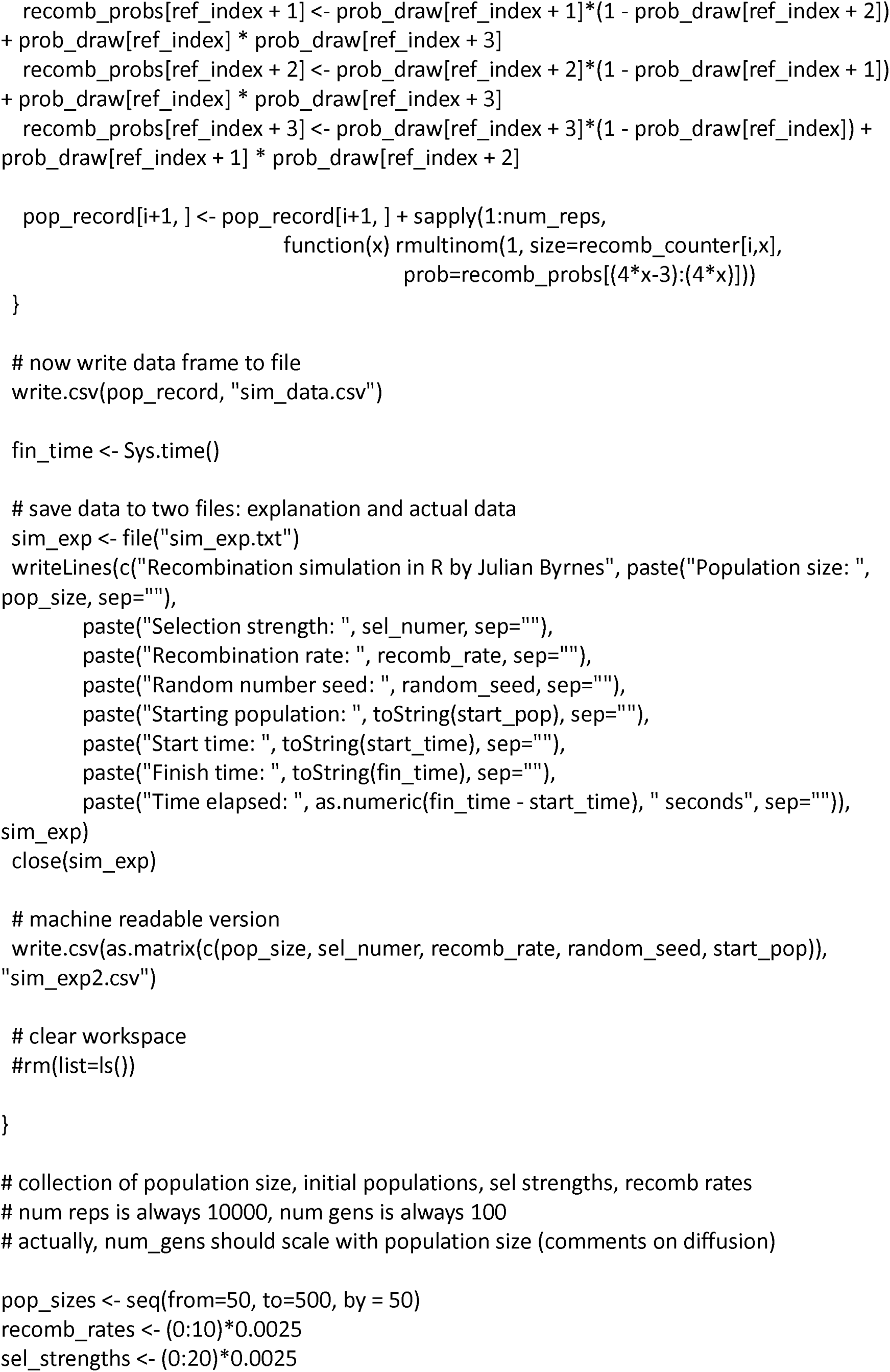
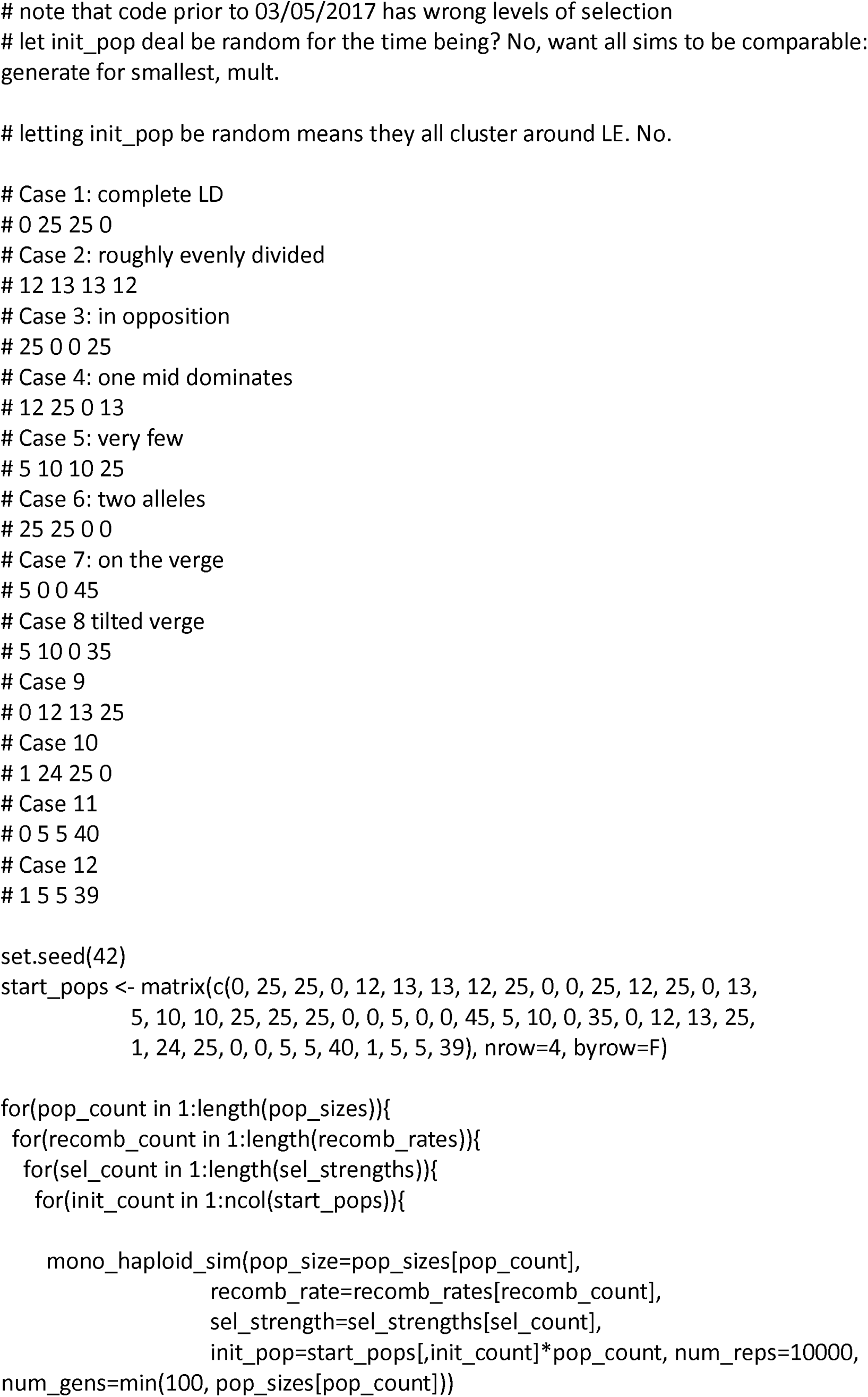
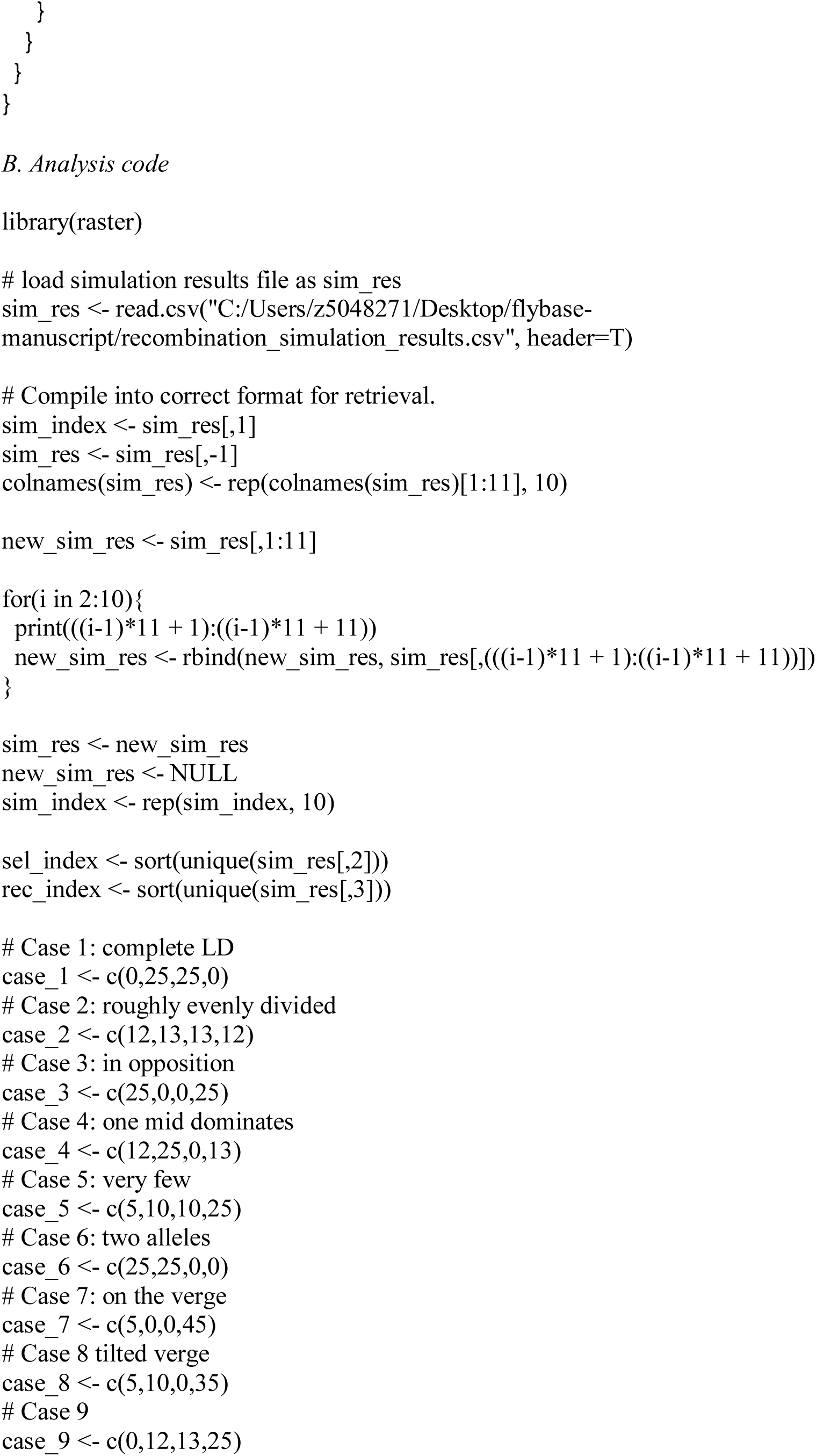
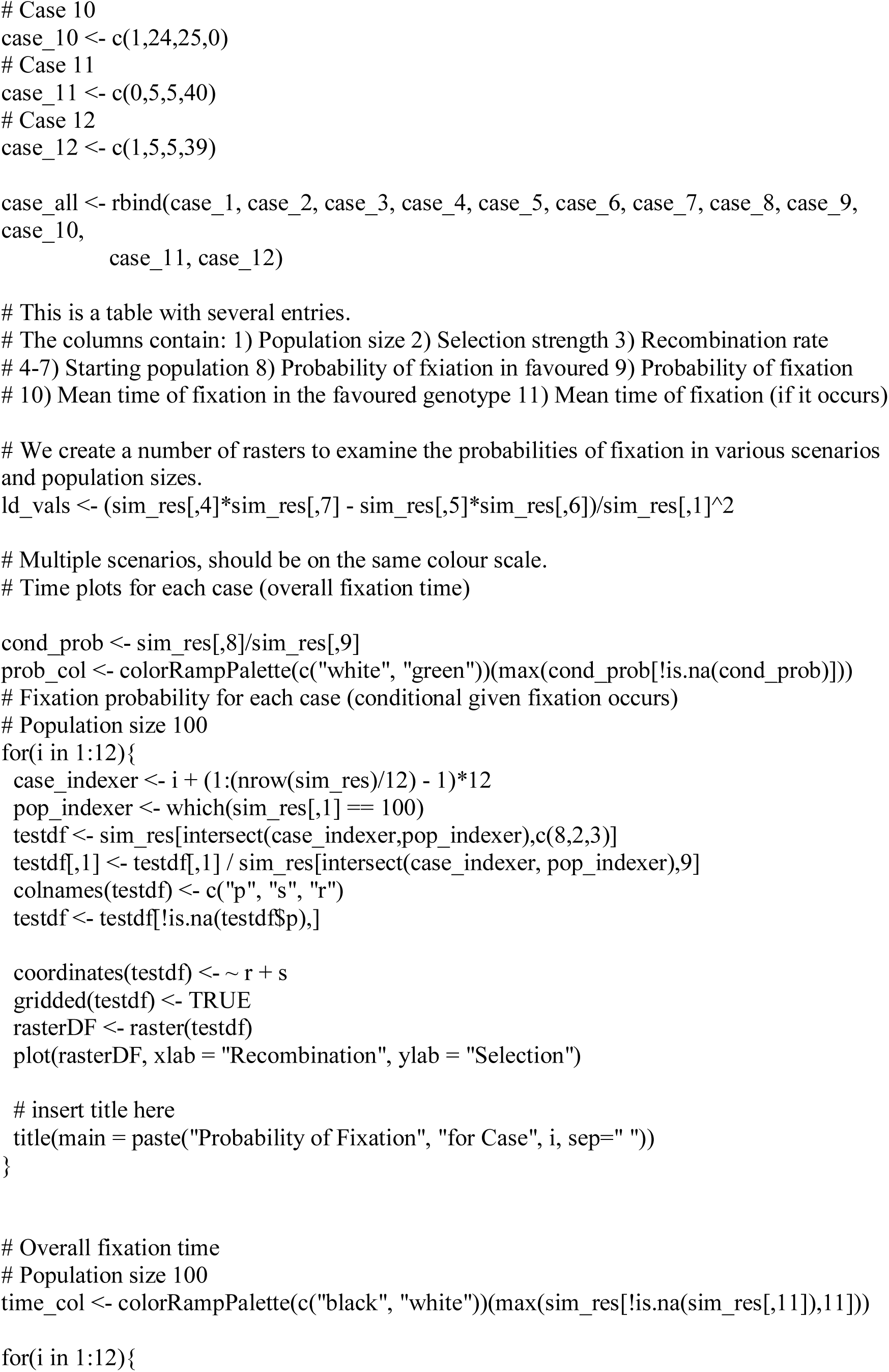
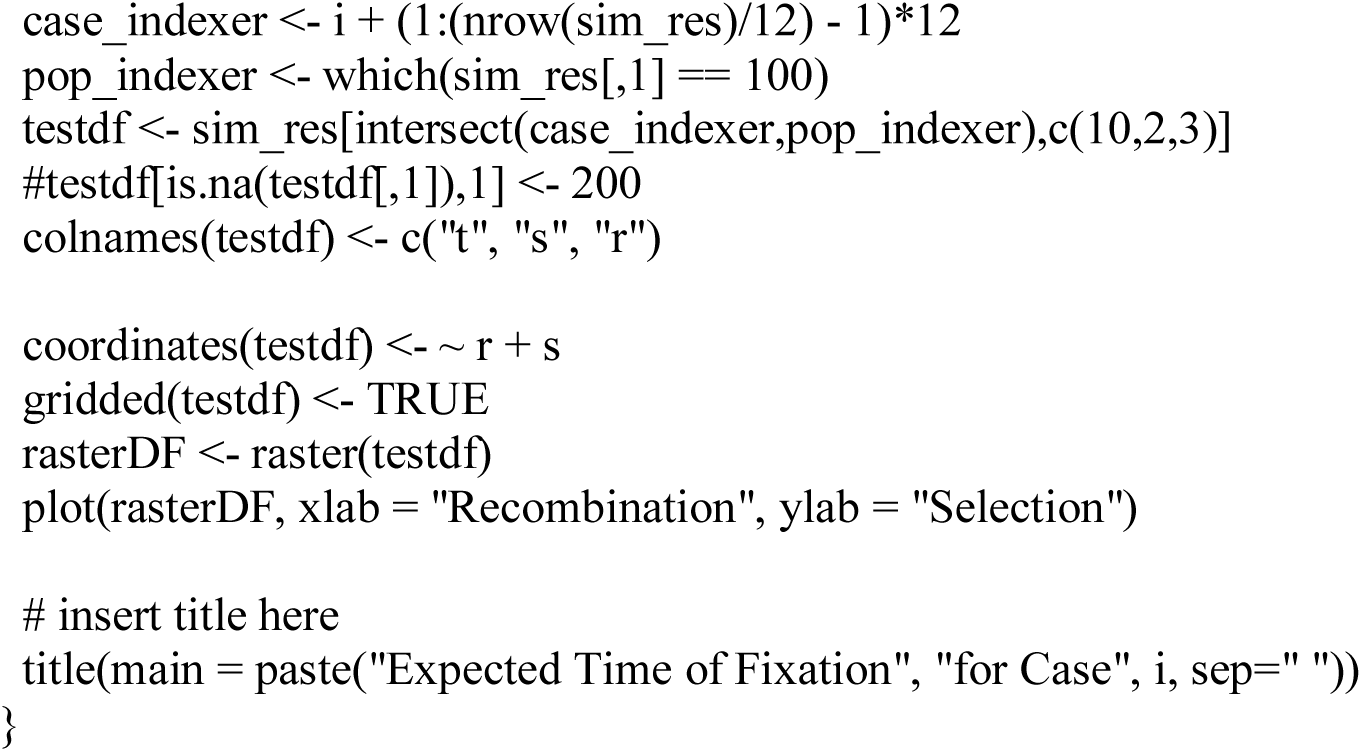

## Figures Supplement

**FIGURE S1:**
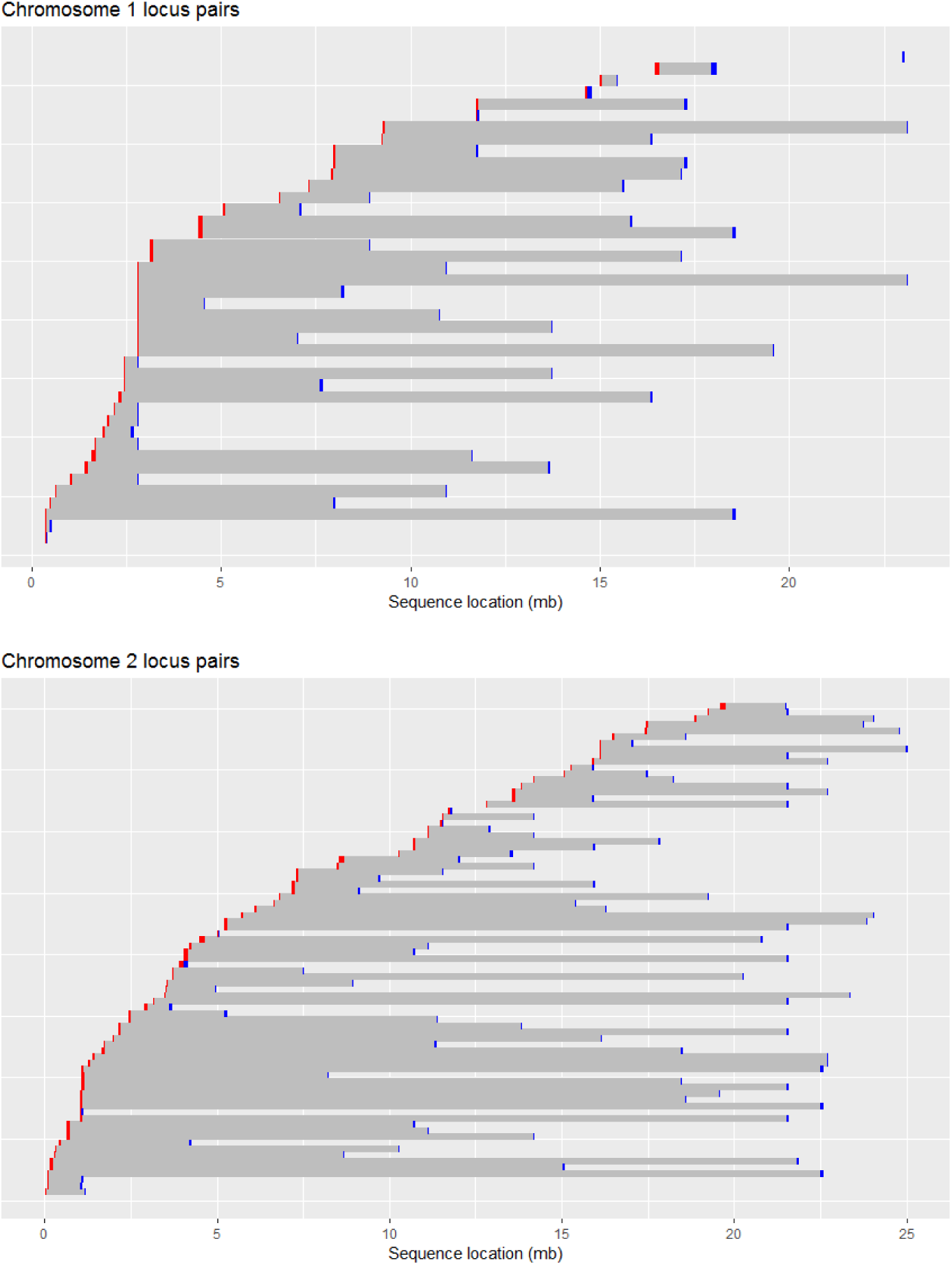
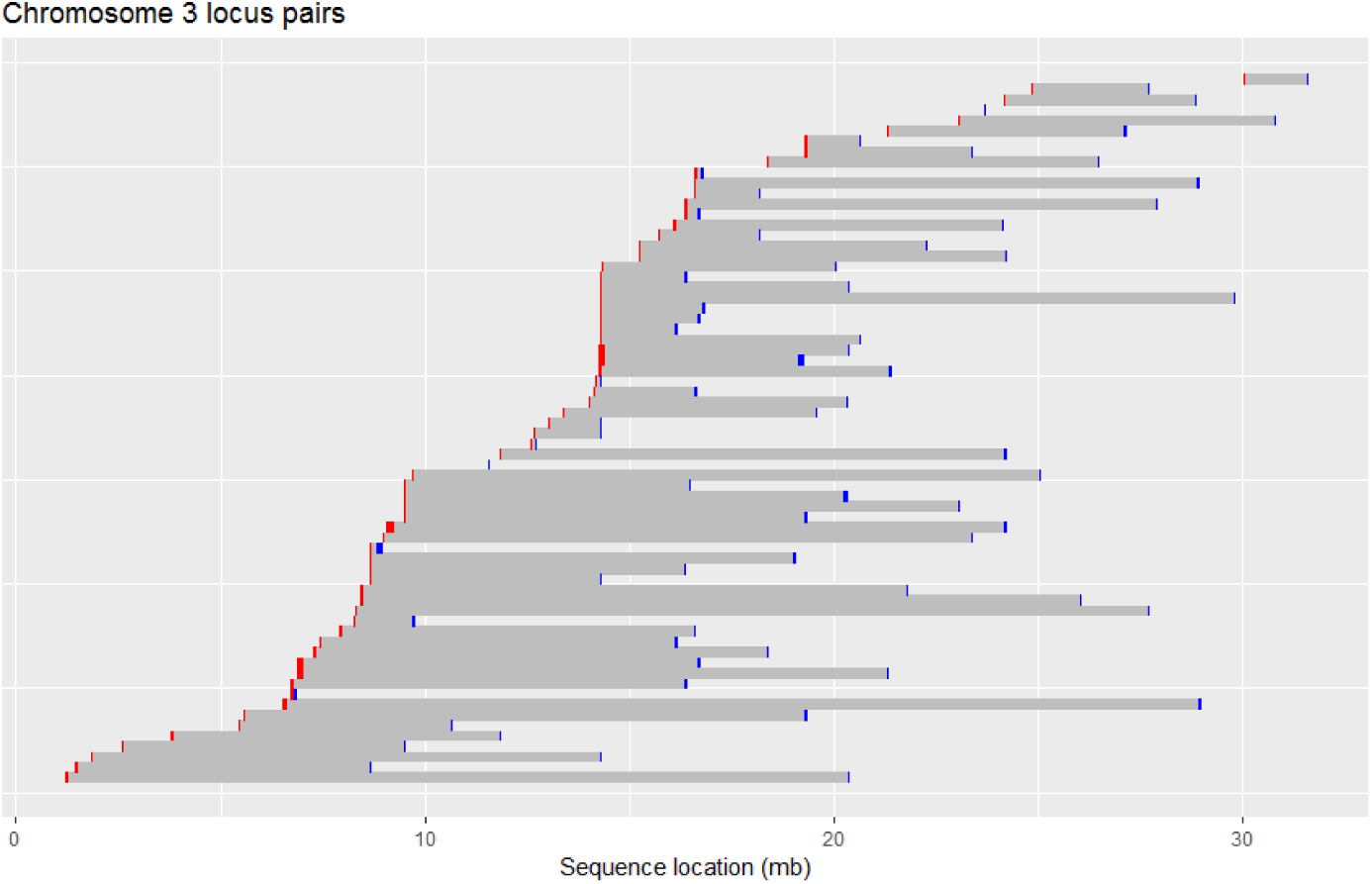
Locations of the locus pairs on each chromosome. Red indicates the ‘leftmost’ locus and blue indicates the ‘rightmost’ locus involved in the particular phenotype. The scale is given in megabases (mb).

**FIGURE S2.**
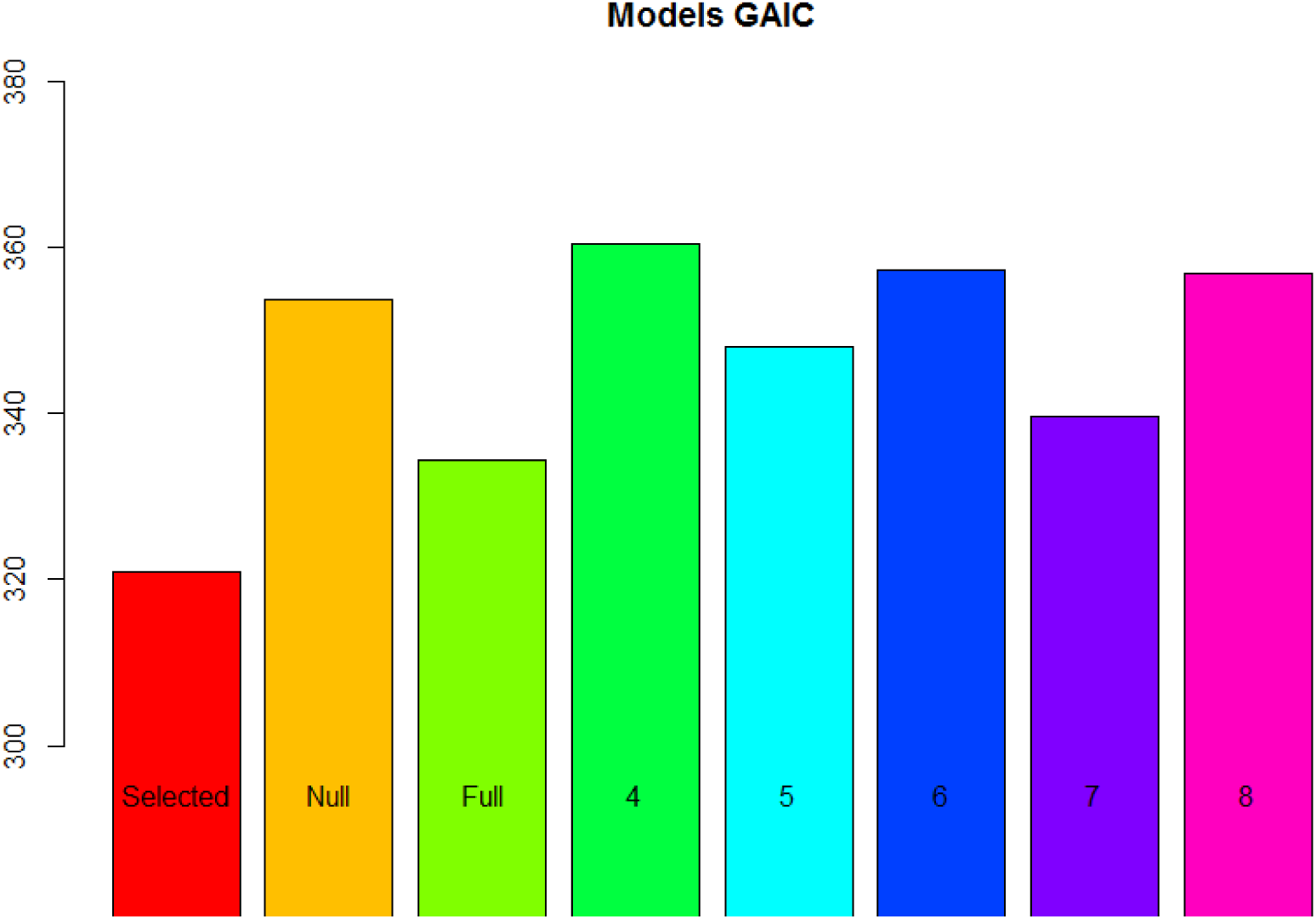
A graphical comparison of the GAICs of the models in [TABLE 6] of the main text.

**FIGURE S3.**
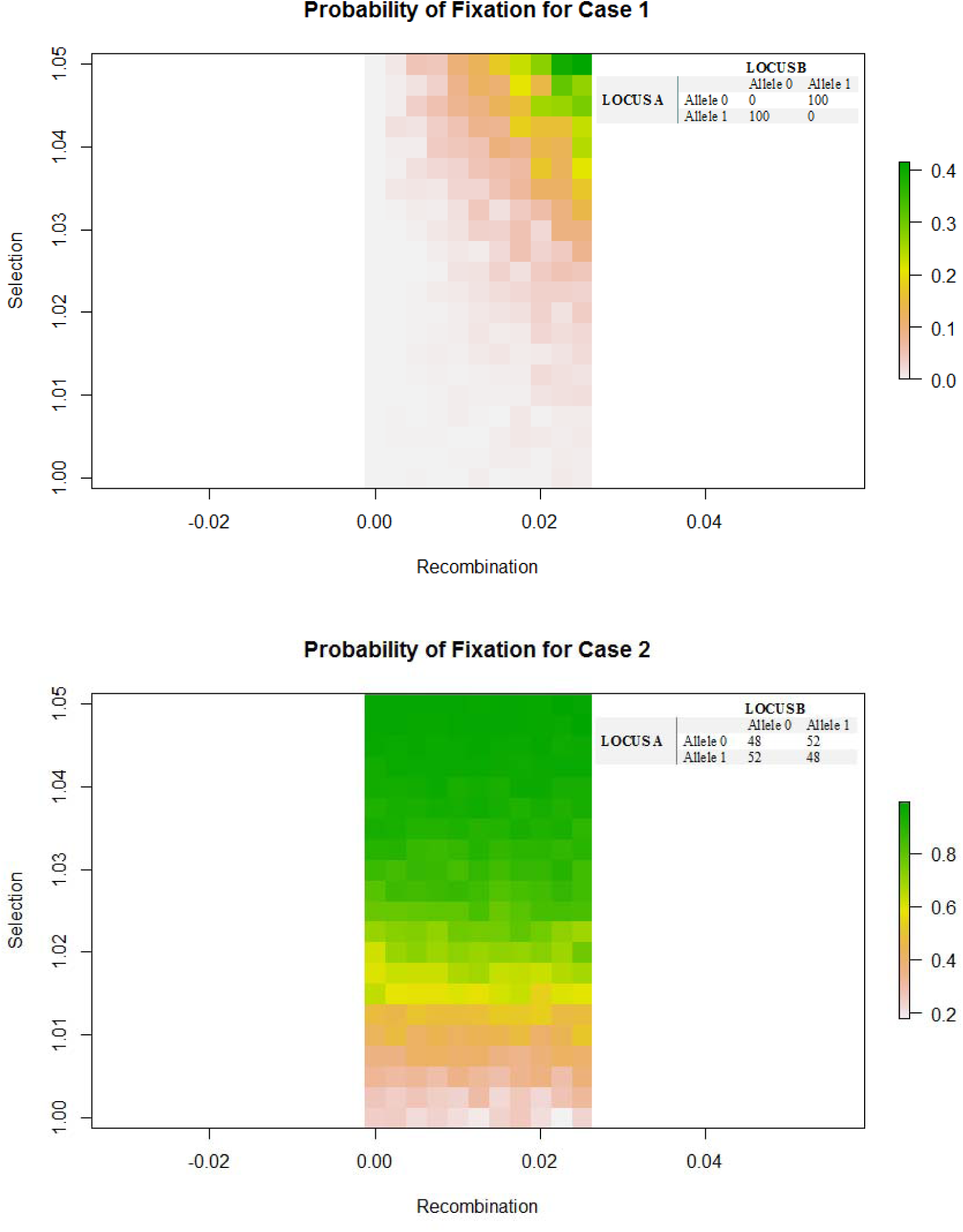
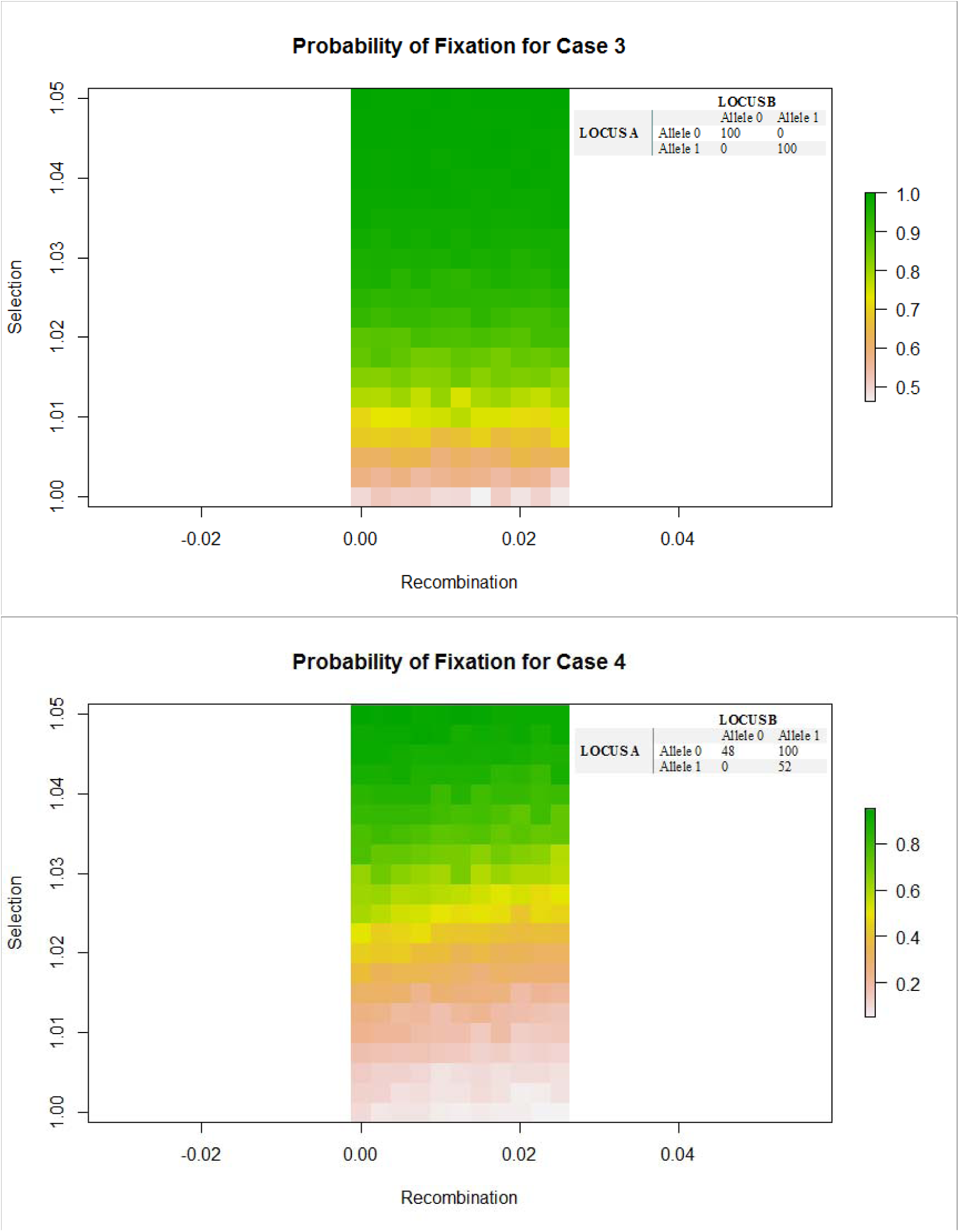
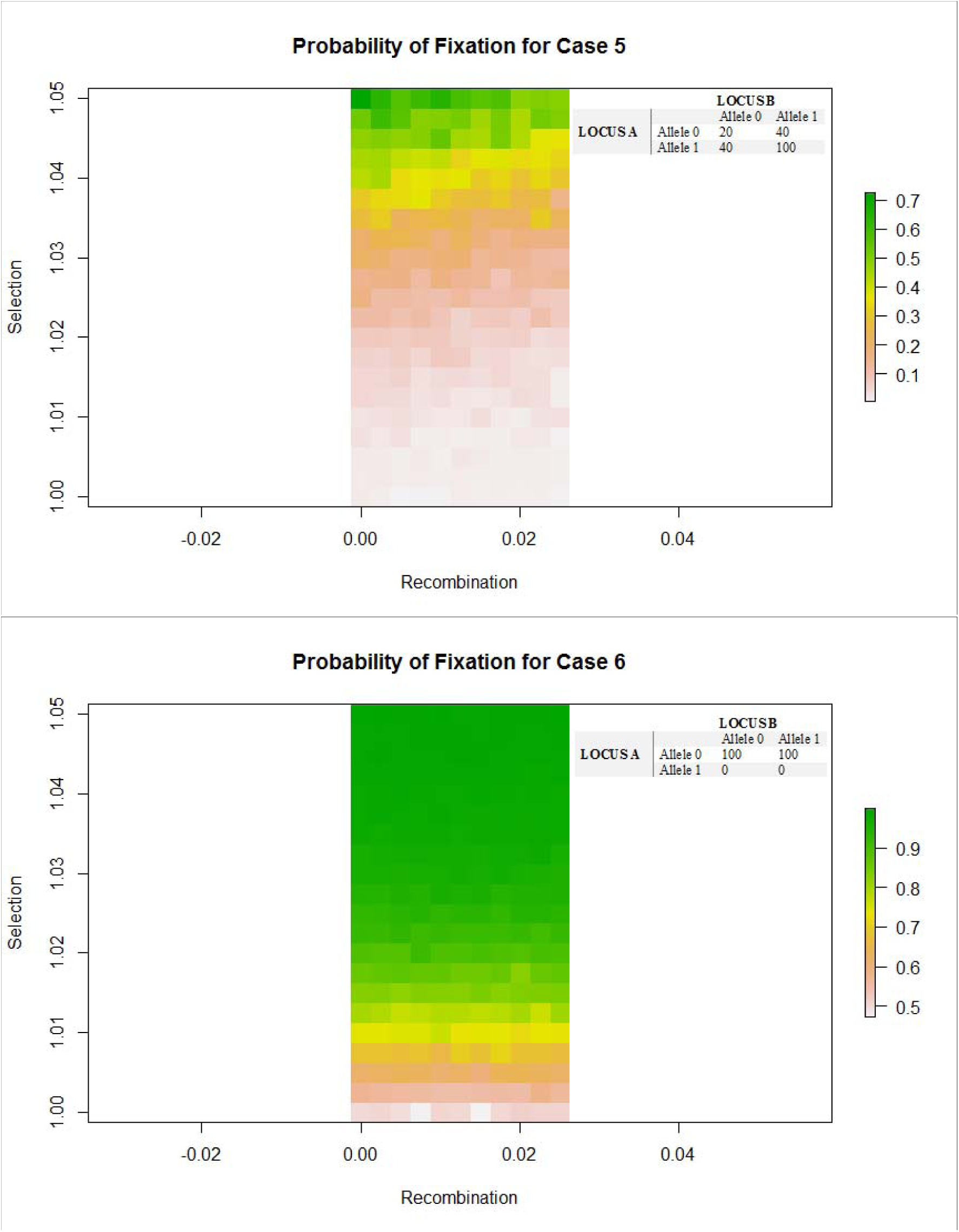
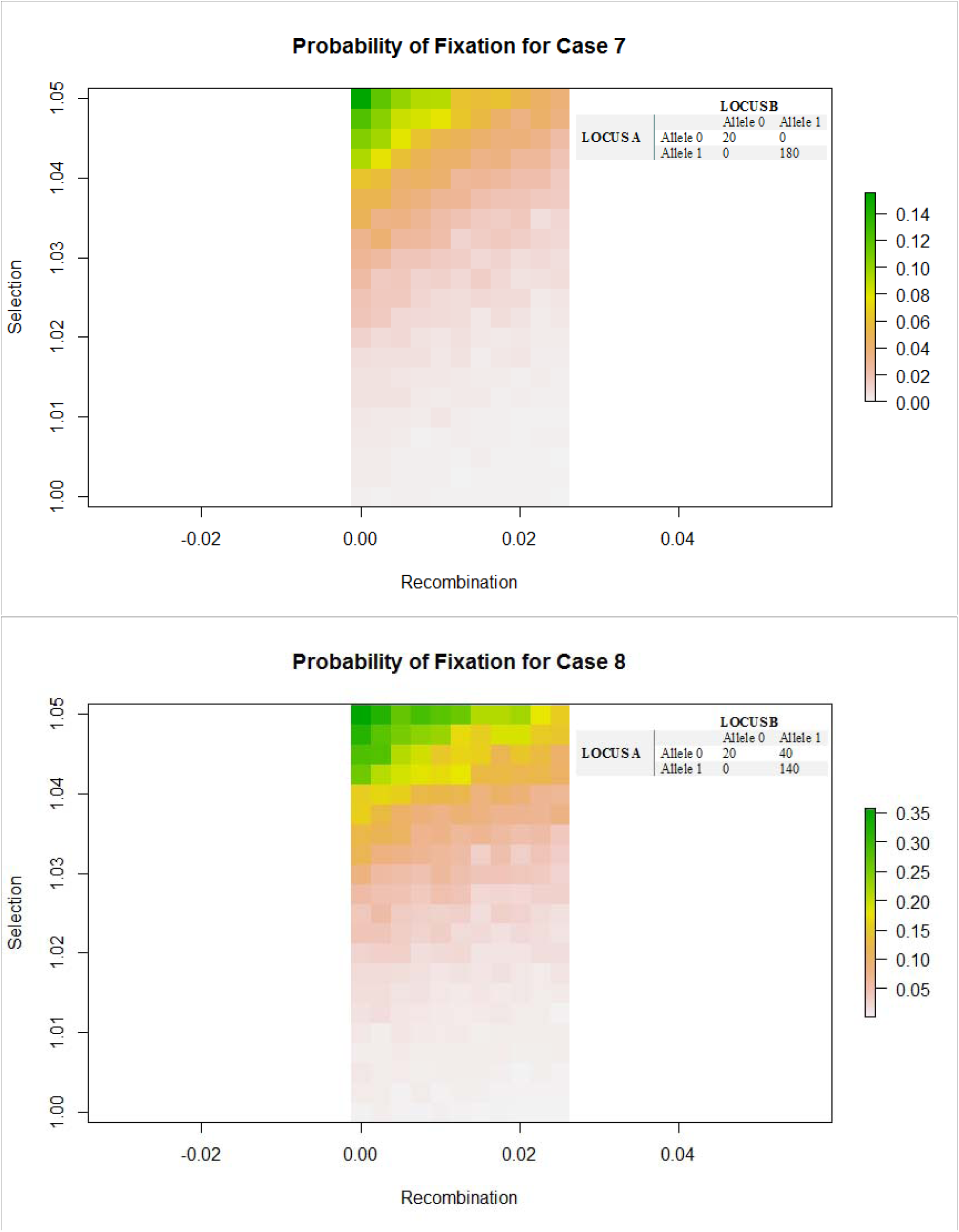
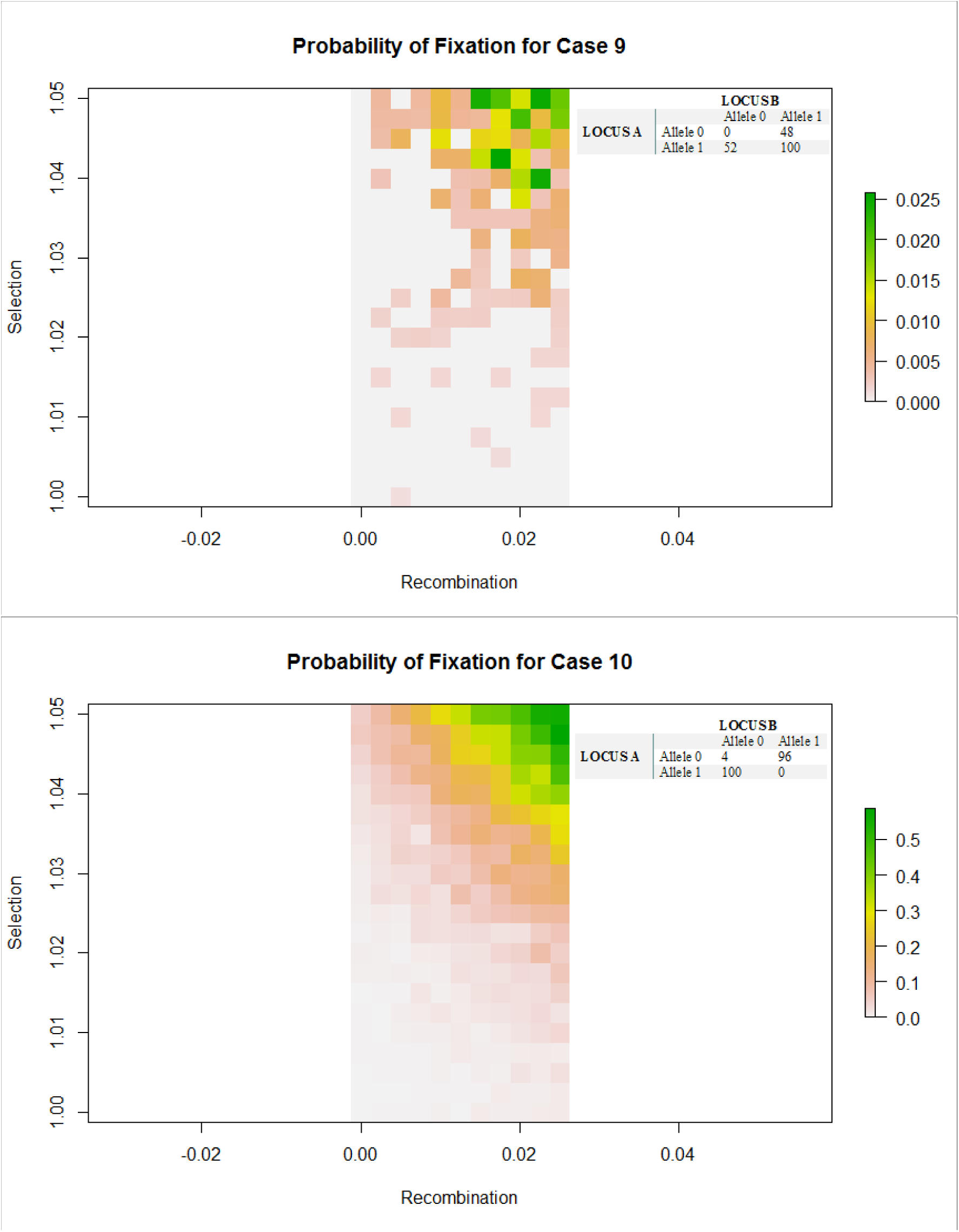
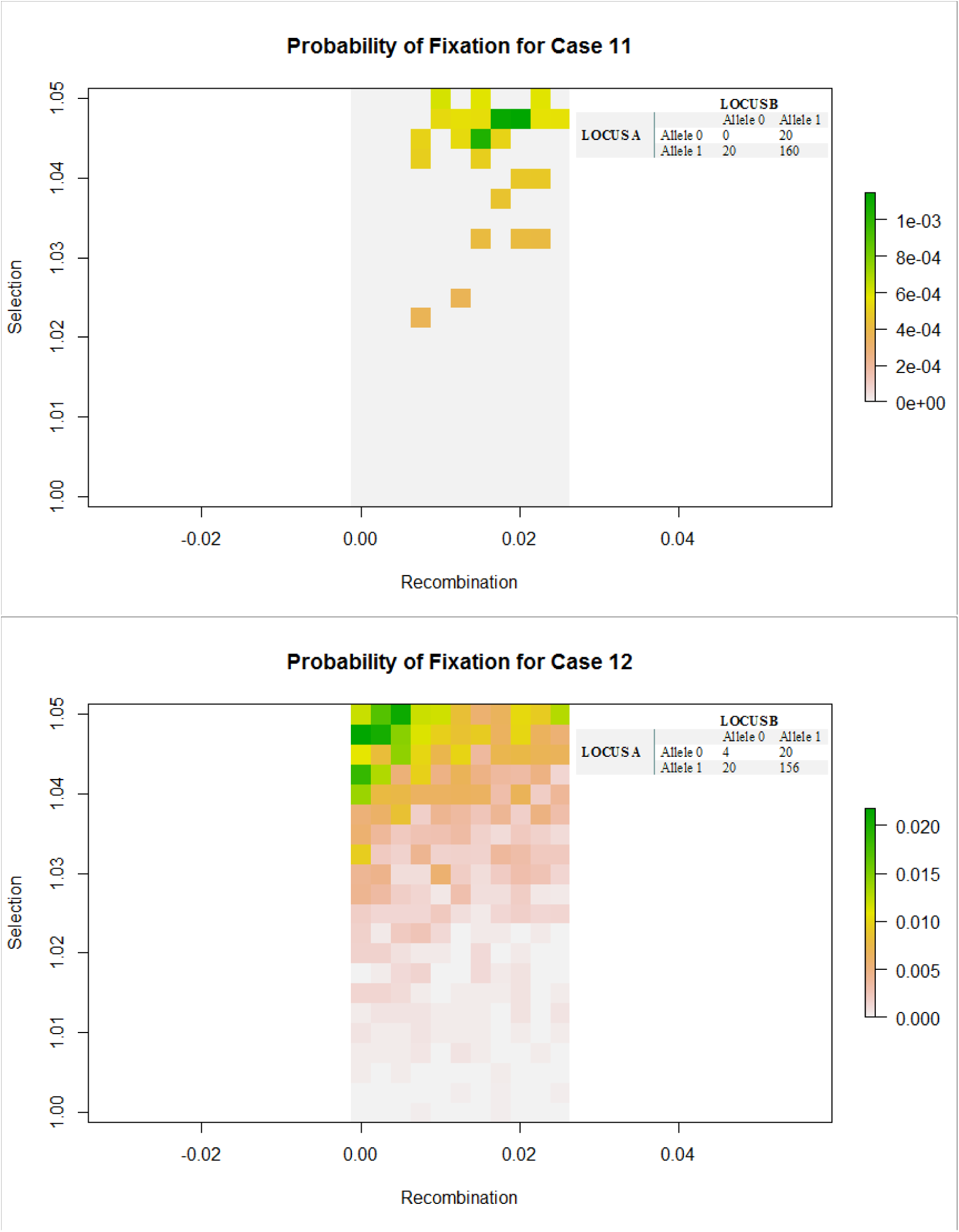
Caption: Rasters of probability of fixation on two axes – recombination and selection. A darker colour indicates a higher probability of fixation (and thus advantage of that recombination and selection combination). The table in the top right corner gives the initial population counts. All graphs are of populations of size 200.

**FIGURE S4:**
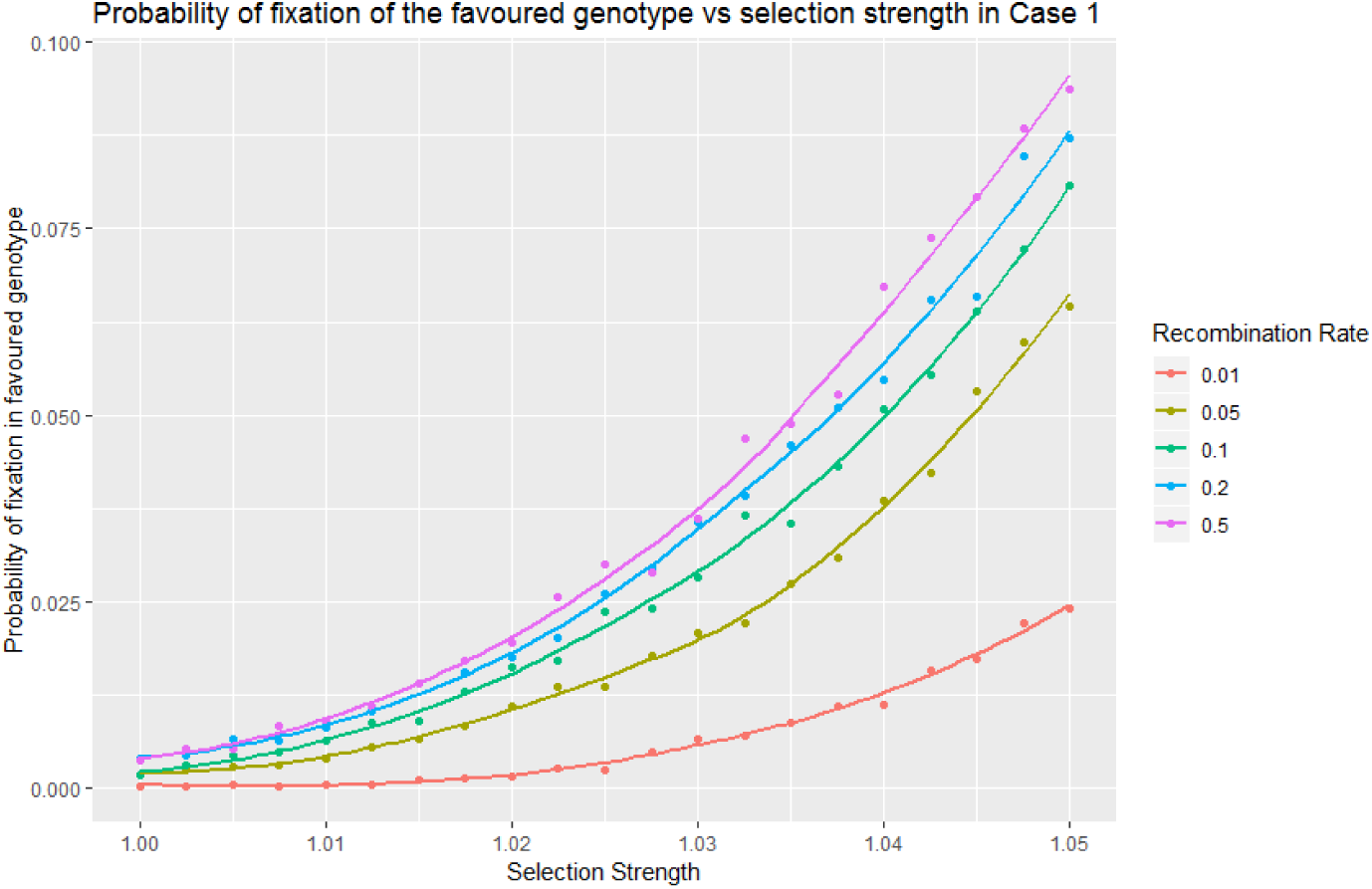
Graph of probability of fixation with higher levels of recombination in [FIGURE S3, Case 1]. These reflect similar patterns to the lower recombination rates.

**FIGURE S5:**
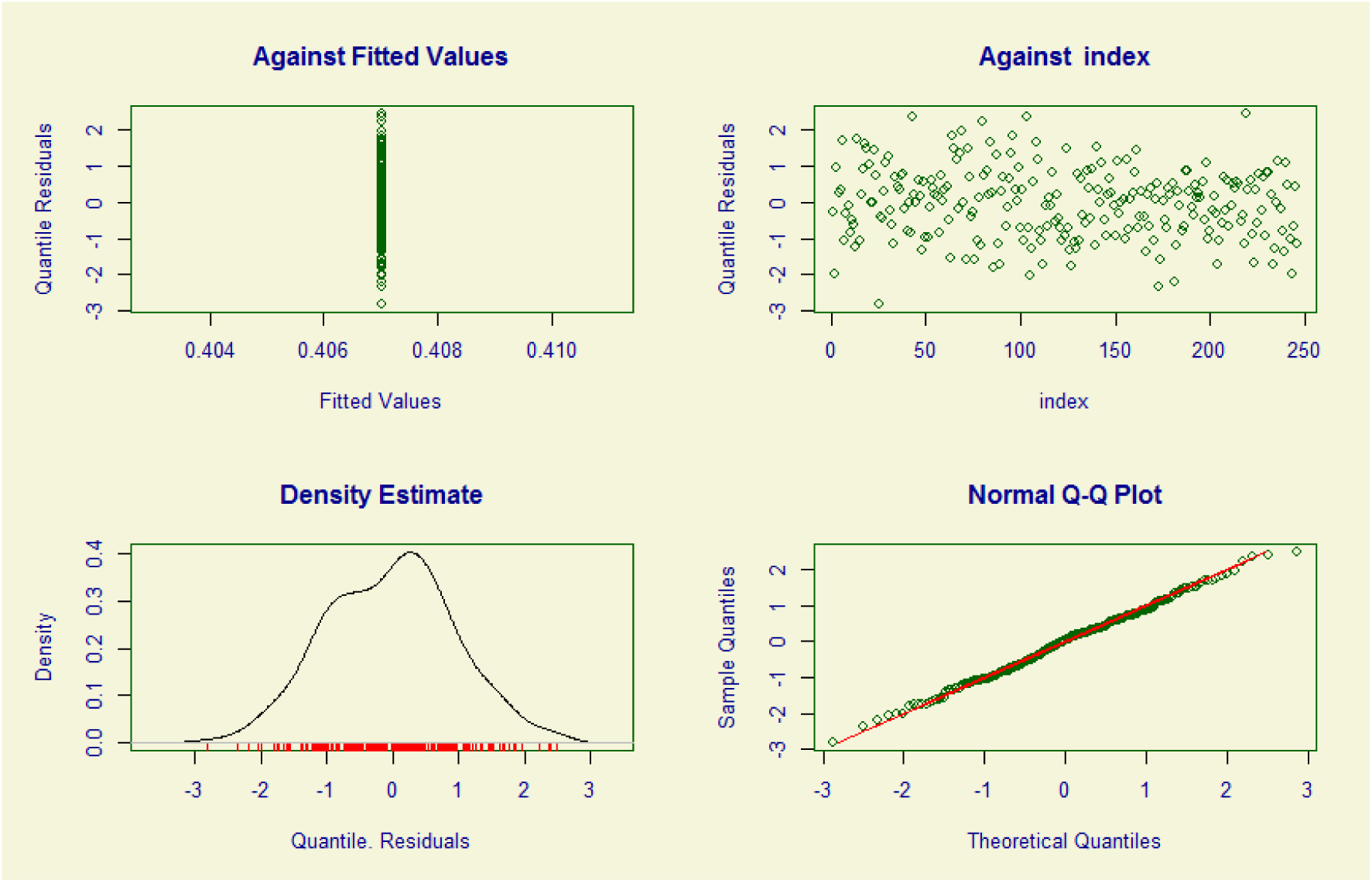
Residual plots for the fitted model. We see that the residuals appear to show no pattern and fit the Q-Q plot very well, indicating a good fit.

